## Supplementary material for "Specialisation of meiotic kinetochores revealed through a synthetic spindle assembly checkpoint strategy": Source Code 1: Supplemental_File_1_kinetochore_synsac.R

```
library(devtools)
library(diann)
library(tidyverse)
library(clipr)
source("/Users/lkoch/Lori MS RStudio/LKfun.R")

# load files ####

pg <- diann_load("final_report.pg_matrix.tsv")
sites_df99 <- diann_load("final_report.phosphosites_99.tsv")

nrow(sites_df99) #4480
nrow(pg) #4761

# shortening sample names

oldnames<-colnames(sites_df99[,7:54])
newnames<-gsub("(.+)_(AM|AM_Lori)_(.+).raw","\\3",oldnames)

# the sample names are the same in the pg file
#colnames(pg)

# rename sample columns to shorter names
# in the pg and sites files
sites_df99<-sites_df99 %>% rename_with(~ newnames, all_of(oldnames))

pg<-pg %>% rename_with(~ newnames, all_of(oldnames))

# for the proteinGroups, we only need to analyse the data in the N columns
# get column numbers of the N intensity cols
columns<-grep("51(.+)N", colnames(pg)) # get intensity column numbers

# plot the 'raw' intensity columns in the pg file
# working with dfs, not se objects here

pg_int_boxplot<-pg[,columns] %>% 
    pivot_longer(cols=all_of(grep("51(.+)N", colnames(pg),value=T)),names_to="sample",values_to="intensity") %>%
    # reorder so metaphase II L comes after metaphase I M
    mutate(sample=factor(sample,levels=c(
        "51K_N-A","51K_N-B","51K_N-C",
        "51K_N-D","51K_N-E","51K_N-F",
        "51M_N-A","51M_N-B","51M_N-C",
        "51M_N-D","51M_N-E","51M_N-F",
        "51L_N-A","51L_N-B","51L_N-C",
        "51L_N-D","51L_N-E","51L_N-F",
        "51N_N-A","51N_N-B","51N_N-C",
        "51N_N-D","51N_N-E","51N_N-F"
    ))) %>%
    ggplot(aes(x=sample,y=log10(intensity)))+
    geom_boxplot()+
    theme(axis.text.x = element_text(angle=90,vjust=0.6))

print_plot(pg_int_boxplot,"pg_int_boxplot.pdf",6,5)

# plot without exp51N-F (no tag rep3) because I do not include it later
print_plot(pg[,columns[1:23]] %>% 
    pivot_longer(cols=colnames(.),names_to="sample",values_to="intensity") %>%
        # reorder so metaphase II L comes after metaphase I M
        mutate(sample=factor(sample,levels=c(
            "51K_N-A","51K_N-B","51K_N-C",
            "51K_N-D","51K_N-E","51K_N-F",
            "51M_N-A","51M_N-B","51M_N-C",
            "51M_N-D","51M_N-E","51M_N-F",
            "51L_N-A","51L_N-B","51L_N-C",
            "51L_N-D","51L_N-E","51L_N-F",
            "51N_N-A","51N_N-B","51N_N-C",
            "51N_N-D","51N_N-E"
        ))) %>%
        mutate(stage=ifelse(grepl("51K",sample),"pro",
                            ifelse(grepl("51M",sample),"MI",
                                   ifelse(grepl("51L",sample),"MII",
                                          "mitoM")))) %>%
        mutate(tag_group=ifelse(grepl("[ABC]",sample),"tag",
                                "no_tag")) %>%
    ggplot(aes(x=sample,y=log10(intensity),fill=interaction(stage,tag_group)))+
    geom_boxplot()+
    theme(axis.text.x = element_text(angle=90,vjust=0.6),
          legend.position="none")+
        labs(title="all protein intensity"),
    "pg_int_boxplot_no51NF.pdf",6,5)

# plots of kinetochore proteins only
kt_protein_pg_int_boxplot<-pg %>% 
    mutate(cat=ifelse(Genes %in% master_kt_df$Gene,"kinetochore","other")) %>%
    filter(cat=="kinetochore") %>%
    select(cat,columns[1:23]) %>%
    pivot_longer(cols=-cat,names_to="sample",values_to="intensity") %>%
    # reorder so metaphase II L comes after metaphase I M
    mutate(sample=factor(sample,levels=c(
        "51K_N-A","51K_N-B","51K_N-C",
        "51K_N-D","51K_N-E","51K_N-F",
        "51M_N-A","51M_N-B","51M_N-C",
        "51M_N-D","51M_N-E","51M_N-F",
        "51L_N-A","51L_N-B","51L_N-C",
        "51L_N-D","51L_N-E","51L_N-F",
        "51N_N-A","51N_N-B","51N_N-C",
        "51N_N-D","51N_N-E"
    ))) %>%
    mutate(stage=ifelse(grepl("51K",sample),"pro",
                        ifelse(grepl("51M",sample),"MI",
                               ifelse(grepl("51L",sample),"MII",
                                      "mitoM")))) %>%
    mutate(tag_group=ifelse(grepl("[ABC]",sample),"tag",
                            "no_tag")) %>%
    ggplot(aes(x=sample,y=log10(intensity),fill=interaction(stage,tag_group)))+
    geom_boxplot()+
    theme(axis.text.x = element_text(angle=90,vjust=0.6),
          legend.position="none")+
    labs(title="kt protein raw int")

print_plot(kt_protein_pg_int_boxplot,"kt_prot_pg_int_boxplot.pdf",6,5)

# colmedians Norm ####
# do this with tags normed to tags 
# and no tags normed to no-tags
# (no cross tag vs tag norm)

pg_tag_cols<-grep("51(.+)N-[ABC]", colnames(pg),value=T)
pg_notag_cols<-grep("51(.+)N-[DEF]", colnames(pg),value=T)
# remove exp51N-F since this rep was too low
pg_notag_cols<-pg_notag_cols[1:11]

colMedians <- function (df, na.rm = T) {
    apply(df, 2, median, na.rm)
}

# calculate target median of tag cols
# then adjust all int to the target med of N cols

colmeds <- colMedians(pg[,pg_tag_cols],na.rm=T)
target_med <- median(colmeds,na.rm=T)
scaling_factors <- as.numeric(target_med/colmeds)
pg[,pg_tag_cols]<-pg[,pg_tag_cols] %>%
    sweep(., 2, scaling_factors, FUN = "*")

# same for no tag cols
colmeds <- colMedians(pg[,pg_notag_cols],na.rm=T)
target_med <- median(colmeds,na.rm=T)
scaling_factors <- as.numeric(target_med/colmeds)
pg[,pg_notag_cols]<-pg[,pg_notag_cols] %>%
    sweep(., 2, scaling_factors, FUN = "*")


# plot intensities after colmedian norm
pg_norm_int_boxplot<-pg[,columns[1:23]] %>%
    pivot_longer(cols=colnames(.),names_to="sample",values_to="intensity") %>%
    # reorder so metaphase II L comes after metaphase I M
    mutate(sample=factor(sample,levels=c(
        "51K_N-A","51K_N-B","51K_N-C",
        "51K_N-D","51K_N-E","51K_N-F",
        "51M_N-A","51M_N-B","51M_N-C",
        "51M_N-D","51M_N-E","51M_N-F",
        "51L_N-A","51L_N-B","51L_N-C",
        "51L_N-D","51L_N-E","51L_N-F",
        "51N_N-A","51N_N-B","51N_N-C",
        "51N_N-D","51N_N-E"
    ))) %>%
    mutate(stage=ifelse(grepl("51K",sample),"pro",
                        ifelse(grepl("51M",sample),"MI",
                               ifelse(grepl("51L",sample),"MII",
                                      "mitoM")))) %>%
    mutate(tag_group=ifelse(grepl("[ABC]",sample),"tag",
                            "no_tag")) %>%
    ggplot(aes(x=sample,y=log10(intensity),fill=interaction(stage,tag_group)))+
    geom_boxplot()+
    theme(axis.text.x = element_text(angle=90,vjust=0.6),
          legend.position="none")+
        labs(title="pg_norm_int")

print_plot(pg_norm_int_boxplot,"pg_norm_int_boxplot.pdf",6,5)

# kinetochore protein intensities after colMedian norm
kt_prot_pg_norm_int_boxplot <- pg %>% 
    mutate(cat=ifelse(Genes %in% master_kt_df$Gene,"kinetochore","other")) %>%
    filter(cat=="kinetochore") %>%
    select(cat,columns[1:23]) %>%
    pivot_longer(cols=-cat,names_to="sample",values_to="intensity") %>%
    # reorder so metaphase II L comes after metaphase I M
    mutate(sample=factor(sample,levels=c(
        "51K_N-A","51K_N-B","51K_N-C",
        "51K_N-D","51K_N-E","51K_N-F",
        "51M_N-A","51M_N-B","51M_N-C",
        "51M_N-D","51M_N-E","51M_N-F",
        "51L_N-A","51L_N-B","51L_N-C",
        "51L_N-D","51L_N-E","51L_N-F",
        "51N_N-A","51N_N-B","51N_N-C",
        "51N_N-D","51N_N-E"
    ))) %>%
    mutate(stage=ifelse(grepl("51K",sample),"pro",
                        ifelse(grepl("51M",sample),"MI",
                               ifelse(grepl("51L",sample),"MII",
                                      "mitoM")))) %>%
    mutate(tag_group=ifelse(grepl("[ABC]",sample),"tag",
                            "no_tag")) %>%
    ggplot(aes(x=sample,y=log10(intensity),fill=interaction(stage,tag_group)))+
        geom_boxplot()+
        theme(axis.text.x = element_text(angle=90,vjust=0.6),
              legend.position="none")+
        labs(title="kt protein colMedian norm")
        
print_plot(kt_prot_pg_norm_int_boxplot,"kt_prot_pg_norm_int_boxplot.pdf",6,5)

# DEP #### 
library(DEP)
# to use DEP functions, need to convert to se obj
# and to do that you need to create these named cols with this function
# DEP needs a 'name' and id column 
pg<-make_unique(pg,"Genes","Protein.Group",delim = ";")

# create modified se which only has 2 replicates for exp51N_F
# # this is because the corresponding PE replicate had too low intensity
exp_design_new<-data.frame(label=grep("51(.+)N",colnames(pg),value=T)[1:23],
                           condition=c(rep("WT_MII",3),rep("no_tag_MII",3),
                                       rep("WT_pro",3),rep("no_tag_pro",3),
                                       rep("WT_MI",3),rep("no_tag_MI",3),
                                       rep("WT_mitoM",3),rep("no_tag_mitoM",2)),
                           replicate=c(rep(c("1","2","3"),7),"1","2"))

# create a summarized Expt object (SE)
pg_se <- make_se(pg, columns[1:23], exp_design_new)

# Filter for proteins that are identified in all replicates of at least one condition 
# choosing this strict filtering since there is such a high identification rate with DIA
pg_se_filt <- filter_missval(pg_se, thr = 0)

# numbers after filtering 
nrow(pg_se) #4761
nrow(pg_se_filt) # 4591

# test differential expression (DEP)

# test DEP 
pg_se_filt_diff <- test_diff(pg_se_filt, type="manual", 
                    test= c("WT_pro_vs_no_tag_pro",
                            "WT_MI_vs_no_tag_MI",
                            "WT_MII_vs_no_tag_MII",
                            "WT_mitoM_vs_no_tag_mitoM",
                            "WT_MII_vs_WT_mitoM",
                            "WT_MII_vs_WT_pro",
                            "WT_MII_vs_WT_MI",
                            "WT_MI_vs_WT_mitoM",
                            "WT_MI_vs_WT_pro",
                            "WT_pro_vs_WT_mitoM"))

# define sig cutoffs
# the pval (alpha) cutoff is the adjusted p-value
# Denote significant proteins based on user defined cutoffs

# set rejections 
dep_pg <- add_rejections_unadj(pg_se_filt_diff, alpha=0.05, log2(2))

# plot PCA
print_plot(plot_pca(dep_pg),"pg_pca.pdf",6,4)

# convert to df obj

dep_pg_df<-get_df_wide(dep_pg)

# VOLCANOS ###

# # DEP volcano function
#plot_volcano(dep_pg,"WT_MI_vs_WT_MII",adjusted=TRUE)

# custom volcanos to get better labels
# load plotting packages
library(ggplot2)
library(plotly)
library(ggrepel)

# now create category column in the MS df
monopolin <- c("CSM1","MAM1","HRR25","LRS4")

dep_pg_df <-dep_pg_df %>%
    mutate(cat=ifelse(name %in% monopolin, "monopolin",
                      ifelse(name %in% master_kt_df$Gene,"kinetochore",
                             ifelse(name %in% spb$Gene,"spindle pole body",
                                    "other"))))

# factor the levels so they always appear in this order

dep_pg_df<-dep_pg_df %>%
    mutate(cat=fct_relevel(cat,c("monopolin","kinetochore","spindle pole body","other")))

## tag vss no tag volcanos
get_volcano_labeled3(dep_pg_df,"WT_pro_vs_no_tag_pro","pro_v_notag_volcano.pdf","pro_v_notag_volcano.html")
get_volcano_labeled3(dep_pg_df,"WT_MI_vs_no_tag_MI","MI_v_notag_volcano.pdf","MI_v_notag_volcano.html")
get_volcano_labeled3(dep_pg_df,"WT_MII_vs_no_tag_MII","MII_v_notag_volcano.pdf","MII_v_notag_volcano.html")
get_volcano_labeled3(dep_pg_df,"WT_mitoM_vs_no_tag_mitoM","mitoM_v_notag_volcano.pdf","mitoM_v_notag_volcano.html")

# Norm to DSN1 level (no tag norm) ####
##  for comparison between different stages 
##  TAG vs TAG only
##  want to do this norm before any other transformations and then basically repeat
##  the above

# what is the value of DSN1 intensity in each column
# still leaving out exp51N rep3

# pg[,columns[1:23]] %>% colnames()

# a matrix of the Dsn1 intensity in
# the columns I want to normalise 5-27
#pg[,5:27] %>% head()
dsn1_df<-pg[pg$name=="DSN1",5:27]

#dsn1_df # 1 row, 23 dsn1 values

# create new duplicate df to hold DSN1 normalised values
# copying matrix "pg" to a new matrix called "dsn1_pg"
dsn1_pg<-pg

# Divide every value in pg matrix columns 5-27
# with the corresponding DSN1 value for that column
# (stored in the dsn1_df 1 row matrix)
# save it into the same columns in the new dsn1_pg matrix
dsn1_pg[,5:27]<-mapply(function(x,y) {x/y},pg[,5:27],  
                       dsn1_df) 
# Dsn1 itself will become 1 in all of these columns
# note the value is a fold change of log2/log2

# distribution of int after dsn1 norm
# bc median of tag conditions is log10 = -2 it means 0.01
# most proteins are 1/100 of dsn1 level in tag conditions
dsn1_norm_int<-dsn1_pg[,columns[1:23]] %>%
    pivot_longer(cols=all_of(colnames(dsn1_pg[,columns[1:23]])),names_to="sample",values_to="intensity") %>%
    # reorder so metaphase II L comes after metaphase I M
    mutate(sample=factor(sample,levels=c(
        "51K_N-A","51K_N-B","51K_N-C",
        "51K_N-D","51K_N-E","51K_N-F",
        "51M_N-A","51M_N-B","51M_N-C",
        "51M_N-D","51M_N-E","51M_N-F",
        "51L_N-A","51L_N-B","51L_N-C",
        "51L_N-D","51L_N-E","51L_N-F",
        "51N_N-A","51N_N-B","51N_N-C",
        "51N_N-D","51N_N-E"
    ))) %>%
    mutate(stage=ifelse(grepl("51K",sample),"pro",
                        ifelse(grepl("51M",sample),"MI",
                               ifelse(grepl("51L",sample),"MII",
                                      "mitoM")))) %>%
    mutate(tag_group=ifelse(grepl("[ABC]",sample),"tag",
                            "no_tag")) %>%
    #pivot_longer(cols=all_of(colnames(dsn1_pg[,columns[1:23]])),names_to="sample",values_to="intensity") %>%
    ggplot(aes(x=sample,y=log10(intensity),fill=interaction(stage,tag_group)))+
    geom_boxplot()+
    theme(axis.text.x = element_text(angle=90,vjust=0.6),
          legend.position="none")+
    labs(title="after normalization to Dsn1p level")


print_plot(dsn1_norm_int,"dsn1_norm_int_plot.pdf",6,5)

# where are kinetochore protein intensities
# before and after norm to Dsn1 levels?
# this will show that the norm makes sense.

kt_dsn1_norm_int<-dsn1_pg %>%
    # filter to only kinetochore proteins
    filter(Genes %in% master_kt_df$Gene) %>%
    select(columns[1:23]) %>%
    pivot_longer(cols=all_of(colnames(dsn1_pg[,columns[1:23]])),names_to="sample",values_to="intensity") %>%
    # reorder so metaphase II L comes after metaphase I M
    mutate(sample=factor(sample,levels=c(
        "51K_N-A","51K_N-B","51K_N-C",
        "51K_N-D","51K_N-E","51K_N-F",
        "51M_N-A","51M_N-B","51M_N-C",
        "51M_N-D","51M_N-E","51M_N-F",
        "51L_N-A","51L_N-B","51L_N-C",
        "51L_N-D","51L_N-E","51L_N-F",
        "51N_N-A","51N_N-B","51N_N-C",
        "51N_N-D","51N_N-E"
    ))) %>%
    mutate(stage=ifelse(grepl("51K",sample),"pro",
                        ifelse(grepl("51M",sample),"MI",
                               ifelse(grepl("51L",sample),"MII",
                                      "mitoM")))) %>%
    mutate(tag_group=ifelse(grepl("[ABC]",sample),"tag",
                            "no_tag")) %>%
    #pivot_longer(cols=all_of(colnames(dsn1_pg[,columns[1:23]])),names_to="sample",values_to="intensity") %>%
    ggplot(aes(x=sample,y=log10(intensity),fill=interaction(stage,tag_group)))+
    geom_boxplot()+
    theme(axis.text.x = element_text(angle=90,vjust=0.6),
          legend.position="none")+
    labs(title="kt prot dsn1 norm")

print_plot(kt_dsn1_norm_int,"kt_dsn1_norm_int_plot.pdf",6,5)

## continuing DEP pipeline with Dsn1-norm data
# to run DEP test_diff on dsn1-norm data, need to convert to se obj
# and do all same steps as above but on this initially norm data

dsn1_se<-make_se(dsn1_pg,columns[1:23],exp_design_new)

# Filter for proteins that are identified in all replicates of at least one condition 
# choosing this strict filtering since there is such a high identification rate with DIA
dsn1_se_filt <- filter_missval(dsn1_se, thr = 0)

# numbers after filtering 
nrow(dsn1_se) #4761
nrow(dsn1_se_filt) # 4591

# test differential expression (DEP)

# test DEP 
dsn1_se_filt_diff <- test_diff(dsn1_se_filt, type="manual", 
                             test= c("WT_MII_vs_WT_mitoM",
                                     "WT_MII_vs_WT_pro",
                                     "WT_MII_vs_WT_MI",
                                     "WT_MI_vs_WT_mitoM",
                                     "WT_MI_vs_WT_pro",
                                     "WT_pro_vs_WT_mitoM"))
# set rejections 
dsn1_dep <- add_rejections_unadj(dsn1_se_filt_diff, alpha=0.05, log2(2))

dsn1_dep_df<-get_df_wide(dsn1_dep)

# Volcanos of dsn1 norm data ####

# now create category column in the MS df

monopolin <- c("CSM1","MAM1","HRR25","LRS4")

dsn1_dep_df <-dsn1_dep_df %>%
    mutate(cat=ifelse(name %in% monopolin, "monopolin",
        ifelse(name %in% master_kt_df$Gene,"kinetochore",
                      ifelse(name %in% spb$Gene,"spindle pole body",
                             "other"))))

# factor the levels so they always appear in this order
dsn1_dep_df<-dsn1_dep_df %>%
    mutate(cat=fct_relevel(cat,c("monopolin","kinetochore","spindle pole body","other")))

# for "dubious" protein names, use yeast systematic names 
# Christos says the Dubious names Wera got from Uniprot
# and I dont find them on SGD so they are not official for yeast

dsn1_dep_df<-dsn1_dep_df %>%
    mutate(name=ifelse(grepl("dubious",name),ID,name)) 
    #filter(grepl("dubious",name)) %>%
    #select(name)

# create dsn1 volcanoes
# only want tag vs tag conditions

# diff stages vs each other
get_volcano_labeled3(dsn1_dep_df,"WT_MI_vs_WT_pro","dsn1_MI_v_pro_volcano.pdf","dsn1_MI_v_pro_volcano.html")
get_volcano_labeled3(dsn1_dep_df,"WT_MII_vs_WT_pro","dsn1_MII_v_pro_volcano.pdf","dsn1_MII_v_pro_volcano.html")
get_volcano_labeled3(dsn1_dep_df,"WT_pro_vs_WT_mitoM","dsn1_pro_v_mitoM_volcano.pdf","dsn1_pro_v_mitoM_volcano.html")
get_volcano_labeled3(dsn1_dep_df,"WT_MII_vs_WT_MI","dsn1_MII_v_MI_volcano.pdf","dsn1_MII_v_MI_volcano.html")
get_volcano_labeled3(dsn1_dep_df,"WT_MI_vs_WT_mitoM","dsn1_MI_v_mitoM_volcano.pdf","dsn1_MI_v_mitoM_volcano.html")
get_volcano_labeled3(dsn1_dep_df,"WT_MII_vs_WT_mitoM","dsn1_MII_v_mitoM_volcano.pdf","dsn1_MII_v_mitoM_volcano.html")

## heatmaps of dsn1 norm prot levels ####

# only want to select tag (NOT NO TAG) cols
dsn1_fc_cols <- factor(c("WT_MII_vs_WT_mitoM_diff",
                         "WT_MII_vs_WT_MI_diff",
                         "WT_MII_vs_WT_pro_diff",
                         "WT_MI_vs_WT_mitoM_diff",
                         "WT_MI_vs_WT_pro_diff",
                         "WT_pro_vs_WT_mitoM_diff"),
                       levels=c("WT_MII_vs_WT_mitoM_diff",
                                "WT_MII_vs_WT_MI_diff",
                                "WT_MII_vs_WT_pro_diff",
                                "WT_MI_vs_WT_mitoM_diff",
                                "WT_MI_vs_WT_pro_diff",
                                "WT_pro_vs_WT_mitoM_diff"))

# top100 proteins ####
# exporting lists of top sigdiff proteins 
# for each stage comparison
# to make excel

# pro v MI
# higher in MI v pro
dsn1_dep_df %>%
    filter(WT_MI_vs_WT_pro_significant==TRUE) %>%
    # filter for direciton of sigdiff
    filter(WT_MI_vs_WT_pro_diff>0) %>%
    arrange(WT_MI_vs_WT_pro_p.val,WT_MI_vs_WT_pro_diff) %>%
    select(name,WT_MI_vs_WT_pro_p.val,WT_MI_vs_WT_pro_diff,cat,First.Protein.Description) %>%
    head(100) %>%
    write_clip()

# higher in pro v MI
dsn1_dep_df %>%
    filter(WT_MI_vs_WT_pro_significant==TRUE) %>%
    # filter for direciton of sigdiff
    filter(WT_MI_vs_WT_pro_diff<0) %>%
    arrange(WT_MI_vs_WT_pro_p.val,WT_MI_vs_WT_pro_diff) %>%
    select(name,WT_MI_vs_WT_pro_p.val,WT_MI_vs_WT_pro_diff,cat,First.Protein.Description) %>%
    head(100) %>%
    write_clip()

# pro v MII
# higher in MII v pro
dsn1_dep_df %>%
    filter(WT_MII_vs_WT_pro_significant==TRUE) %>%
    # filter for direciton of sigdiff
    filter(WT_MII_vs_WT_pro_diff>0) %>%
    arrange(WT_MII_vs_WT_pro_p.val,WT_MII_vs_WT_pro_diff) %>%
    select(name,WT_MII_vs_WT_pro_p.val,WT_MII_vs_WT_pro_diff,cat,First.Protein.Description) %>%
    head(100) %>%
    write_clip()

# higher in pro v MII
dsn1_dep_df %>%
    filter(WT_MII_vs_WT_pro_significant==TRUE) %>%
    # filter for direciton of sigdiff
    filter(WT_MII_vs_WT_pro_diff<0) %>%
    arrange(WT_MII_vs_WT_pro_p.val,WT_MII_vs_WT_pro_diff) %>%
    select(name,WT_MII_vs_WT_pro_p.val,WT_MII_vs_WT_pro_diff,cat,First.Protein.Description) %>%
    head(100) %>%
    write_clip()

# pro v mitoM - WT_pro_vs_WT_mitoM
# higher in pro v mitoM
dsn1_dep_df %>%
    filter(WT_pro_vs_WT_mitoM_significant==TRUE) %>%
    # filter for direciton of sigdiff
    filter(WT_pro_vs_WT_mitoM_diff>0) %>%
    arrange(WT_pro_vs_WT_mitoM_p.val,WT_pro_vs_WT_mitoM_diff) %>%
    select(name,WT_pro_vs_WT_mitoM_p.val,WT_pro_vs_WT_mitoM_diff,cat,First.Protein.Description) %>%
    head(100) %>%
    write_clip()

# higher in mitoM v pro
dsn1_dep_df %>%
    filter(WT_pro_vs_WT_mitoM_significant==TRUE) %>%
    # filter for direciton of sigdiff
    filter(WT_pro_vs_WT_mitoM_diff<0) %>%
    arrange(WT_pro_vs_WT_mitoM_p.val,WT_pro_vs_WT_mitoM_diff) %>%
    select(name,WT_pro_vs_WT_mitoM_p.val,WT_pro_vs_WT_mitoM_diff,cat,First.Protein.Description) %>%
    head(100) %>%
    write_clip()

# MI v MII
# higher in MI vs MII
dsn1_dep_df %>%
    filter(WT_MII_vs_WT_MI_significant==TRUE) %>%
    # filter for direciton of sigdiff
    filter(WT_MII_vs_WT_MI_diff<0) %>%
    arrange(WT_MII_vs_WT_MI_p.val,WT_MII_vs_WT_MI_diff) %>%
    select(name,WT_MII_vs_WT_MI_p.val,WT_MII_vs_WT_MI_diff,cat,First.Protein.Description) %>%
    head(100) %>%
    write_clip()

# higher in MII vs MI
dsn1_dep_df %>%
    filter(WT_MII_vs_WT_MI_significant==TRUE) %>%
    # filter for direciton of sigdiff
    filter(WT_MII_vs_WT_MI_diff>0) %>%
    arrange(WT_MII_vs_WT_MI_p.val,WT_MII_vs_WT_MI_diff) %>%
    select(name,WT_MII_vs_WT_MI_p.val,WT_MII_vs_WT_MI_diff,cat,First.Protein.Description) %>%
    head(100) %>%
    write_clip()

# MI v mitoM
# # higher in MI v mitoM
dsn1_dep_df %>%
    filter(WT_MI_vs_WT_mitoM_significant==TRUE) %>%
    # filter for direciton of sigdiff
    filter(WT_MI_vs_WT_mitoM_diff>0) %>%
    arrange(WT_MI_vs_WT_mitoM_p.val,WT_MI_vs_WT_mitoM_diff) %>%
    select(name,WT_MI_vs_WT_mitoM_p.val,WT_MI_vs_WT_mitoM_diff,cat,First.Protein.Description) %>%
    head(100) %>%
    write_clip()

# # higher in mitoM v MI
dsn1_dep_df %>%
    filter(WT_MI_vs_WT_mitoM_significant==TRUE) %>%
    # filter for direciton of sigdiff
    filter(WT_MI_vs_WT_mitoM_diff<0) %>%
    arrange(WT_MI_vs_WT_mitoM_p.val,WT_MI_vs_WT_mitoM_diff) %>%
    select(name,WT_MI_vs_WT_mitoM_p.val,WT_MI_vs_WT_mitoM_diff,cat,First.Protein.Description) %>%
    head(100) %>%
    write_clip()

# MII v mitoM
# # higher in MII v mitoM
dsn1_dep_df %>%
    filter(WT_MII_vs_WT_mitoM_significant==TRUE) %>%
    # filter for direciton of sigdiff
    filter(WT_MII_vs_WT_mitoM_diff>0) %>%
    arrange(WT_MII_vs_WT_mitoM_p.val,WT_MII_vs_WT_mitoM_diff) %>%
    select(name,WT_MII_vs_WT_mitoM_p.val,WT_MII_vs_WT_mitoM_diff,cat,First.Protein.Description) %>%
    head(100) %>%
    write_clip()

# # higher in mitoM v MII
dsn1_dep_df %>%
    filter(WT_MII_vs_WT_mitoM_significant==TRUE) %>%
    # filter for direciton of sigdiff
    filter(WT_MII_vs_WT_mitoM_diff<0) %>%
    arrange(WT_MII_vs_WT_mitoM_p.val,WT_MII_vs_WT_mitoM_diff) %>%
    select(name,WT_MII_vs_WT_mitoM_p.val,WT_MII_vs_WT_mitoM_diff,cat,First.Protein.Description) %>%
    head(100) %>%
    write_clip()

### new kinetochore list heatmaps ####

# break up master_kt_df (n=79)
# into shorter lists

# length(cbf1_3) # 5 # really 4
# length(inner_kt) # 18 # really 17
# length(KMN) #10 # really 10
# length(dam1) #10 # really 10
# length(accessory) #19 # really 15
# length(maps) #8 # really 7
# length(cpc_sac) # really 9


cbf1_3<-c("NDC10","CEP3","CTF13","SKP1","CBF1")

inner_kt<-c("MIF2","CSE4","CTF19","OKP1","MCM21","AME1","CHL4","CNN1",
            "WIP1","MHF1","MHF2","MCM16","CTF3","MCM22","IML3","NKP1",
            "NKP2","YBP2")

KMN<-c("MTW1","DSN1","NNF1","NSL1","NDC80","NUF2","SPC24","SPC25","SPC105",
       "KRE28")

dam1<-c("ASK1","DAD1","DAD2","DAD3","DAD4","DAM1","DUO1","SPC19","SPC34",
        "HSK3")

accessory<-c("SLK19","PAT1","FIN1","MAM1","CSM1","HRR25","LRS4","SPO13",
             "SLX8","GLC7","SPT4","CRM1","RTS1","SGO1","RIO1","ULS1",
             "PLC1","CBF2","TPD3")

maps<-c("KIP1","KIP3","CIN8","KAR3","BIK1","STU1","STU2","KAR9")

cpc_sac<-c("IPL1","SLI15","NBL1","BIR1","MAD1","MAD2","BUB1","BUB3","MPS1")

#  tag/notag KT prot heatmap ####
# # remember this MUST come from non-Dsn1 normalised DF
# 
tag_notag_fcs<-grep("_diff",colnames(dep_pg_df),value=T) %>%
    grep("no_tag",.,value=T)

tag_notag_sig_cols<-grep("_significant",colnames(dep_pg_df),value=T) %>%
    grep("no_tag",.,value=T)

tag_notag_pval_cols<-grep("_p.val",colnames(dep_pg_df),value=T) %>%
    grep("no_tag",.,value=T)
# 
# # factor contrast levels so they are in the stage order I want...
# 

pg_kt_tag_notag_heatmap<-dep_pg_df %>%
    filter(cat=="kinetochore") %>%
    select(tag_notag_fcs, name, tag_notag_sig_cols,tag_notag_pval_cols) %>%
    # filter out proteins that have NA fc in all these contrasts
    filter(if_any(-c(name,tag_notag_sig_cols,tag_notag_pval_cols),~!is.na(.x))) %>%
    pivot_longer(-c(name,tag_notag_sig_cols,tag_notag_pval_cols),names_to="sample",values_to="fc") %>%
    ### the line below is important for plotting in the right order
    #select(name,sample,fc) %>% head()
    mutate(sample2=gsub("_diff","",sample)) %>%
    mutate(sample2=factor(sample2,levels=c("WT_pro_vs_no_tag_pro",
                                         "WT_MI_vs_no_tag_MI",
                                         "WT_MII_vs_no_tag_MII",
                                         "WT_mitoM_vs_no_tag_mitoM"),ordered=T)) %>%
    mutate(name=fct_reorder2(name,dplyr::desc(sample2),dplyr::desc(fc),.na_rm=F)) %>%
    # mutate(sig_label=ifelse(WT_MII_vs_WT_MI_significant==TRUE,
    #                         ifelse(WT_MII_vs_WT_MI_p.val<0.001,"***",
    #                                ifelse(WT_MII_vs_WT_MI_p.val<0.01,"**",
    #                                       ifelse(WT_MII_vs_WT_MI_p.val<0.05,"*",NA))),NA)) %>%
    ggplot(aes(x=sample2,y=name,fill=fc))+
    geom_tile(color="black")+
    scale_fill_distiller(palette="RdBu",limits=c(-9,9))+
    #geom_text(aes(),size=3)+
    theme_minimal()+
    theme(axis.text.x = element_text(angle=90,vjust=0.5))+
    labs(x="",y="",fill="log2fc",title="tag v no tag") +
    coord_flip()


print_plot(pg_kt_tag_notag_heatmap,"pg_kt_tag_notag_heatmap.pdf",12,4)

# 
# # KT prot heatmap all samples ####
# # instead of showing fold changes, simply plot the abundance
# # of the proteins in all 8 samples (4x tag and 4x no tag)

# variable to hold all sample columns
pg_colnames<-colnames(dep_pg_df)[2:24]

pg_kt_all_samples_heatmap<-dep_pg_df %>%
    filter(cat=="kinetochore") %>%
    select(name, pg_colnames) %>%
    # filter out proteins that have NA fc in all these contrasts
    filter(if_any(-c(name),~!is.na(.x))) %>%
    pivot_longer(-c(name),names_to="sample",values_to="log2int") %>%
    ### the line below is important for plotting in the right order
    #select(name,sample,fc) %>% head()
   # mutate(sample2=gsub("_diff","",sample)) %>%
    #mutate(sample=factor(sample,levels=c("WT_pro_vs_no_tag_pro",
                                           # "WT_MI_vs_no_tag_MI",
                                           # "WT_MII_vs_no_tag_MII",
                                           # "WT_mitoM_vs_no_tag_mitoM"),ordered=T)) %>%
    mutate(sample=factor(sample, levels=c("WT_mitoM_1","WT_mitoM_2","WT_mitoM_3",
                                          "WT_MII_1","WT_MII_2","WT_MII_3",
                                          "WT_MI_1","WT_MI_2","WT_MI_3",
                                          "WT_pro_1","WT_pro_2","WT_pro_3",
                                          "no_tag_mitoM_1","no_tag_mitoM_2","no_tag_mitoM_3",
                                          "no_tag_MII_1","no_tag_MII_2","no_tag_MII_3",
                                          "no_tag_MI_1","no_tag_MI_2","no_tag_MI_3",
                                          "no_tag_pro_1","no_tag_pro_2","no_tag_pro_3"))) %>%
    mutate(name=fct_reorder2(name,dplyr::desc(sample),dplyr::desc(log2int),.na_rm=F)) %>%
    mutate(int=2^log2int) %>%
    # mutate(sig_label=ifelse(WT_MII_vs_WT_MI_significant==TRUE,
    #                         ifelse(WT_MII_vs_WT_MI_p.val<0.001,"***",
    #                                ifelse(WT_MII_vs_WT_MI_p.val<0.01,"**",
    #                                       ifelse(WT_MII_vs_WT_MI_p.val<0.05,"*",NA))),NA)) %>%
    ggplot(aes(x=sample,y=name,fill=log10(int)))+
    geom_tile(color="black")+
    #scale_fill_distiller(palette="PuOr")+
    scale_fill_gradient(low="white",high="darkgreen")+
    #geom_text(aes(),size=3)+
    theme_minimal()+
    theme(axis.text.x = element_text(angle=90,vjust=0.5))+
    labs(x="",y="",fill="log10int",title="kinetochore proteins") +
    coord_flip()

print_plot(pg_kt_all_samples_heatmap,"pg_kt_all_samples_heatmap.pdf",12,4)

# BEGINNING PHOSPHO ####
# PE norm to PG ####
# 
sites_df99 <- sites_df99 %>%
    mutate(gene_name_site=paste0(Gene.Names,"_",Site))

pe_99<-make_unique(sites_df99,"gene_name_site","Sequence",delim = ";")

# column NUMBERs of PE cols
pe_cols<-grep("51(.+)PE",colnames(pe_99))

# convert zeros to NAs so they are ignored by data transformations

pe_99[,pe_cols]<-pe_99[,pe_cols] %>%
    apply(.,2,function(x) ifelse(x==0,NA,x))

# 'raw' intensity boxplot
pe_99_int_boxplot<-pe_99[,pe_cols[1:23]] %>%
    pivot_longer(cols=all_of(grep("51(.+)PE", colnames(pe_99),value=T)[1:23]),names_to="sample",values_to="intensity") %>%
    ggplot(aes(x=sample,y=log10(intensity)))+
    geom_boxplot()+
    theme(axis.text.x = element_text(angle=90,vjust=0.6))

print_plot(pe_99_int_boxplot,"pe_99_int_boxplot.pdf",6,5)

# hopefully there are more sites in teh PE vs N cols...
pg_sites_99_int_boxplot<-pe_99[,grep("51(.+)N",colnames(pe_99),value=T)[1:23]] %>%
    pivot_longer(cols=all_of(grep("51(.+)N",colnames(pe_99),value=T)[1:23]),names_to="sample",values_to="intensity") %>%
    ggplot(aes(x=sample,y=log10(intensity)))+
    geom_boxplot()+
    theme(axis.text.x = element_text(angle=90,vjust=0.6))+
    labs(title="phospho-site int in N cols")

print_plot(pg_sites_99_int_boxplot,"pg_sites_99_int_boxplot.pdf",6,5)

# TAGs should be norm SEPARATELY from no tags

pe_tag_cols<-grep("51(.+)PE-[ABC]",colnames(pe_99),value=T)
pe_notag_cols<-grep("51(.+)PE-[DEF]",colnames(pe_99),value=T)
pe_notag_cols<-pe_notag_cols[1:11] # remove bad sample
pe_tag_cols<-pe_tag_cols[c(1:9,11:12)] # remove bad sample

# colmedian normalize tag samples
colmeds <- colMedians(pe_99[,pe_tag_cols],na.rm=T)
target_med <- median(colmeds,na.rm=T)
scaling_factors <- as.numeric(target_med/colmeds)
pe_99[,pe_tag_cols]<-pe_99[,pe_tag_cols] %>%
    sweep(., 2, scaling_factors, FUN = "*")

# colmedian normalise no tag samples
colmeds <- colMedians(pe_99[,pe_notag_cols],na.rm=T)
target_med <- median(colmeds,na.rm=T)
scaling_factors <- as.numeric(target_med/colmeds)
pe_99[,pe_notag_cols]<-pe_99[,pe_notag_cols] %>%
    sweep(., 2, scaling_factors, FUN = "*")

# # plot norm
# # 
# # colnames(pe_99[,pe_cols[1:23]])
# # colnames(pe_99[,pe_notag_cols])

pe_norm_int_boxplot<-pe_99[,pe_cols[1:23]] %>%
    pivot_longer(cols=all_of(colnames(.)),names_to="sample",values_to="intensity") %>%
    ggplot(aes(x=sample,y=log10(intensity)))+
    geom_boxplot()+
    theme(axis.text.x = element_text(angle=90,vjust=0.6))

# you can see that the no-tag cols have higher medians now...
print_plot(pe_norm_int_boxplot,"pe_norm_int_boxplot.pdf",6,5)

# the pe_99 df has changed zeros to NAs
# and only has relevant columns

# # normalize each phospho-site to the corresponding protein 
# 
# # colnames(pg)
# # head(pg$Protein.Group)
# # head(pe_99$Protein)
# # looks like in pg Protein.Group (with splitstrp)
# # in pe_99 use Protein
# 
strsplit.extract <- function (x, split, index) {
    x <- as.character(x)
    x[x == ""] <- ";"
    x <- sapply(strsplit(x, split, fixed = TRUE), "[[", index)
    return(x)
}

pg$first.Protein.Group<-strsplit.extract(pg$Protein.Group,";",1)

phos.stoic.temp.df <- pg[match(pe_99$Protein, pg$first.Protein.Group), columns[c(1:18,20:23)]]

#this divides the intensity of the phosphos over the protg
#normalized intensity then multiplies by 1000 (to get easier number)
#This creates "stoichiometric" phos. site changes...

# create duplicate df to hold prot norm pe

pe_pg_99<-pe_99

# perform norm to protein level

pe_pg_99[,pe_cols[c(1:18,20:23)]] <- mapply(function(x,y) {x/y},pe_99[,pe_cols[c(1:18,20:23)]],  phos.stoic.temp.df) * 1000

# plot distribution after prot norm

pe_pg_99_int_boxplot<-pe_pg_99[,pe_cols[c(1:18,20:23)]] %>%
    pivot_longer(cols=all_of(grep("51(.+)PE", colnames(pe_pg_99),value=T)[c(1:18,20:23)]),names_to="sample",values_to="intensity") %>%
    ggplot(aes(x=sample,y=log10(intensity)))+
    geom_boxplot()+
    theme(axis.text.x = element_text(angle=90,vjust=0.6))

print_plot(pe_pg_99_int_boxplot,"pe_pg_99_int_boxplot.pdf",6,5)

# create a summarized Expt object (SE)

# # create a design matrix
# # ignoring the 3 rep of exp51N
pe_exp_design_new<-data.frame(label=grep("51(.+)PE",colnames(pe_99),value=T)[c(1:18,20:23)],
                           condition=c(rep("WT_MII",3),rep("no_tag_MII",3),
                                       rep("WT_pro",3),rep("no_tag_pro",3),
                                       rep("WT_MI",3),rep("no_tag_MI",3),
                                       rep("WT_mitoM",2),rep("no_tag_mitoM",2)),
                           replicate=c(rep(c("1","2","3"),6),"1","2","1","2"))

pe_pg_99_se <- make_se(pe_pg_99, pe_cols[c(1:18,20:23)], pe_exp_design_new)

# filter for detections in 3/3 of at least 1 condition

pe_pg_99_se_filt <- filter_missval(pe_pg_99_se, thr = 0)

# how many sites after filtering
length(pe_pg_99_se) # 4480
length(pe_pg_99_se_filt) # 2211

plot_numbers(pe_pg_99_se_filt)

# tests DEP
pe_pg_99_se_filt_diff<-test_diff(pe_pg_99_se_filt,type="manual",
                                   test= c("WT_pro_vs_no_tag_pro",
                                           "WT_MI_vs_no_tag_MI",
                                           "WT_MII_vs_no_tag_MII",
                                           "WT_mitoM_vs_no_tag_mitoM",
                                           "WT_pro_vs_WT_MI",
                                           "WT_pro_vs_WT_MII",
                                           "WT_pro_vs_WT_mitoM",
                                           "WT_MI_vs_WT_MII",
                                           "WT_MI_vs_WT_mitoM",
                                           "WT_MII_vs_WT_mitoM"))

dep_pe_pg_99_filt_diff <- add_rejections_unadj(pe_pg_99_se_filt_diff, alpha = 0.05, lfc = log2(2))

# convert to df
dep_pe_pg_99_filt_df<-get_df_wide(dep_pe_pg_99_filt_diff)

# PE_PG plots ####

# how many sites were quantified (in 3/3 of one condition)
# in each condition

# remember now the int cols are cols 2:24 named WT_MII_1
# etc
#
pe_int_colnames<-colnames(dep_pe_pg_99_filt_df[,2:23])

sites_per_sample_bar<-dep_pe_pg_99_filt_df %>%
    select(all_of(pe_int_colnames)) %>%
    mutate(across(all_of(pe_int_colnames),function(x) sum(x>0,na.rm=T))) %>%
    head(1) %>%
    pivot_longer(cols=all_of(pe_int_colnames),values_to="number",names_to="sample") %>%
    ggplot(aes(x=fct_inorder(sample),y=number))+
    geom_bar(aes(fill=sample),stat="identity")+
    geom_text(aes(label=number),vjust=0.5,hjust=1.3)+
    theme_classic()+
    theme(legend.position="none")+
    coord_flip()+
    scale_y_continuous(expand=expansion(mult=c(0,0.1)))+ # 10% more plot area after max value
    labs(y="",x="sample",title="sites per sample")

print_plot(sites_per_sample_bar,"sites_per_sample_bar.pdf",7.5,5)

# how many sites are sig in each contrast
pe_sig_cols<-grep("_significant",colnames(dep_pe_pg_99_filt_df),value=T)

pe_sites_per_contrast<-dep_pe_pg_99_filt_df %>%
    filter(significant==T) %>%
    select(all_of(pe_sig_cols)) %>%
    apply(.,2,function(x) sum(x,na.rm=T)) %>%
    data.frame(number=.) %>%
    cbind(contrast=rownames(.)) %>%
    mutate(contrast=gsub("_significant","",contrast)) %>%
    ggplot(aes(x=contrast,y=number,fill=contrast))+
    geom_bar(stat="identity")+
    theme_classic()+
    theme(axis.text.x = element_text(angle=90,vjust=0.5,hjust=1),
          legend.position="none")+
    scale_y_continuous(expand=expansion(mult=c(0,0.2)))+ # 20% more plot area after max value
    labs(title="number of sig sites per contrast")

print_plot(pe_sites_per_contrast,"pe_sites_per_contrast.pdf",5,5)

## OLD MOTIFs at each stage ####
## ALL SITES FROM IP #### (NOT JUST KT PROTEIN SITES)

library(ggseqlogo)

pe_col_names<-colnames(dep_pe_pg_99_filt_df[,2:23])

MII_tag_cols<-pe_col_names[1:3]
MII_notag_cols<-pe_col_names[4:6]
pro_tag_cols<-pe_col_names[7:9]
pro_notag_cols<-pe_col_names[10:12]
MI_tag_cols<-pe_col_names[13:15]
MI_notag_cols<-pe_col_names[16:18]
mitoM_tag_cols<-pe_col_names[19:20]
mitoM_notag_cols<-pe_col_names[21:22]

# make a df holding these sums to plot
MII_tag_num<-dep_pe_pg_99_filt_df %>% 
    # select only sites detected in 3/3 reps of that stage
    filter_at(MII_tag_cols,~ .x > 0) %>%
    nrow()

MII_notag_num<-dep_pe_pg_99_filt_df %>% 
    # select only sites detected in 3/3 reps of that stage
    filter_at(MII_notag_cols,~ .x > 0) %>%
    nrow()

MI_tag_num<-dep_pe_pg_99_filt_df %>% 
    # select only sites detected in 3/3 reps of that stage
    filter_at(MI_tag_cols,~ .x > 0) %>%
    nrow()

MI_notag_num<-dep_pe_pg_99_filt_df %>% 
    # select only sites detected in 3/3 reps of that stage
    filter_at(MI_notag_cols,~ .x > 0) %>%
    nrow()

pro_tag_num<-dep_pe_pg_99_filt_df %>% 
    # select only sites detected in 3/3 reps of that stage
    filter_at(pro_tag_cols,~ .x > 0) %>%
    nrow()

pro_notag_num<-dep_pe_pg_99_filt_df %>% 
    # select only sites detected in 3/3 reps of that stage
    filter_at(pro_notag_cols,~ .x > 0) %>%
    nrow()

mitoM_tag_num<-dep_pe_pg_99_filt_df %>% 
    # select only sites detected in 3/3 reps of that stage
    filter_at(mitoM_tag_cols,~ .x > 0) %>%
    nrow()

mitoM_notag_num<-dep_pe_pg_99_filt_df %>% 
    # select only sites detected in 3/3 reps of that stage
    filter_at(mitoM_notag_cols,~ .x > 0) %>%
    nrow()

pe_num_df<-data.frame(number=c(MII_tag_num,MII_notag_num,
                       MI_tag_num,MI_notag_num,
                       pro_tag_num,pro_notag_num,
                       mitoM_tag_num,mitoM_notag_num),
              sample = c("MII_tag","MII_notag",
                            "MI_tag","MI_notag",
                            "pro_tag","pro_notag",
                            "mitoM_tag","mitoM_notag"))

pe_sites_3reps<-pe_num_df %>%
    ggplot(aes(x=sample,y=number))+
    geom_bar(aes(fill=sample),stat="identity")+
    geom_text(aes(label=number),vjust=0.5,hjust=1.3)+
    theme_classic()+
    theme(legend.position="none")+
    coord_flip()+
    scale_y_continuous(expand=expansion(mult=c(0,0.1)))+ # 10% more plot area after max value
    labs(y="",x="sample",title="sites in 3/3 reps")

print_plot(pe_sites_3reps,"pe_sites_3reps.pdf",5,3)

# PROTEIN VENN ####

# how many proteins are shared or unique in each sample:
# pro 51K -- 
# MI 51M -- 
# MII 51L -- 
# mitoM 51N -- 

# cutoff all replicates detection (at least 1) vs 0/3
pg_colnames<-colnames(dep_pg_df)[2:24]
pro_cols<-pg_colnames[7:9]
MI_cols <- pg_colnames[13:15]
MII_cols <-pg_colnames[1:3]
mitoM_cols <- pg_colnames[19:21]

dep_pg_df<-dep_pg_df %>%
    mutate(detect_MI=apply(.[,MI_cols],1,function(x) !any(is.na(x)))) %>%
    mutate(detect_MII=apply(.[,MII_cols],1,function(x) !any(is.na(x)))) %>%
    mutate(detect_pro=apply(.[,pro_cols],1,function(x) !any(is.na(x)))) %>%
    mutate(detect_mitoM=apply(.[,mitoM_cols],1,function(x) !any(is.na(x)))) %>%
    mutate(detect_MI_notag=apply(.[,pg_colnames[16:18]],1,function(x) !any(is.na(x)))) %>%
    mutate(detect_MII_notag=apply(.[,pg_colnames[4:6]],1,function(x) !any(is.na(x)))) %>%
    mutate(detect_pro_notag=apply(.[,pg_colnames[10:12]],1,function(x) !any(is.na(x)))) %>%
    mutate(detect_mitoM_notag=apply(.[,pg_colnames[21:22]],1,function(x) !any(is.na(x))))

# export protein name lists for http://www.interactivenn.net/

# MI
dep_pg_df %>%
    filter(detect_MI==TRUE) %>%
    select(name) %>%
    write_clip()

# MII
dep_pg_df %>%
    filter(detect_MII==TRUE) %>%
    select(name) %>%
    write_clip()

# pro
dep_pg_df %>%
    filter(detect_pro==TRUE) %>%
    select(name) %>%
    write_clip()

# mitoM
dep_pg_df %>%
    filter(detect_mitoM==TRUE) %>%
    select(name) %>%
    write_clip()

# PE detection overlaps ####
# PHOS VENN ####
# 
# cutoff ALL replicates detection
# pe_int_colnames
# pe_int_colnames[21:22]
pe_pro_cols<-pe_int_colnames[7:9]
pe_MI_cols <- pe_int_colnames[13:15]
pe_MII_cols <-pe_int_colnames[1:3]
pe_mitoM_cols <- pe_int_colnames[19:20]

# detect_MI means it was detected in 3/3 reps of MI
dep_pe_pg_99_filt_df <- dep_pe_pg_99_filt_df  %>%
    mutate(detect_MI=apply(.[,pe_MI_cols],1,function(x) !any(is.na(x)))) %>%
    mutate(detect_MII=apply(.[,pe_MII_cols],1,function(x) !any(is.na(x)))) %>%
    mutate(detect_pro=apply(.[,pe_pro_cols],1,function(x) !any(is.na(x)))) %>%
    mutate(detect_mitoM=apply(.[,pe_mitoM_cols],1,function(x) !any(is.na(x)))) %>%
    mutate(detect_MI_notag=apply(.[,pe_int_colnames[16:18]],1,function(x) !any(is.na(x)))) %>%
    mutate(detect_MII_notag=apply(.[,pe_int_colnames[4:6]],1,function(x) !any(is.na(x)))) %>%
    mutate(detect_pro_notag=apply(.[,pe_int_colnames[10:12]],1,function(x) !any(is.na(x)))) %>%
    mutate(detect_mitoM_notag=apply(.[,pe_int_colnames[21:22]],1,function(x) !any(is.na(x))))


# export protein name lists for http://www.interactivenn.net/

# MI
dep_pe_pg_99_filt_df %>%
    filter(detect_MI==TRUE) %>%
    select(name) %>%
    write_clip()

# MII
dep_pe_pg_99_filt_df %>%
    filter(detect_MII==TRUE) %>%
    select(name) %>%
    write_clip()

# pro
dep_pe_pg_99_filt_df %>%
    filter(detect_pro==TRUE) %>%
    select(name) %>%
    write_clip()

# mitoM
dep_pe_pg_99_filt_df %>%
    filter(detect_mitoM==TRUE) %>%
    select(name) %>%
    write_clip()


# pe kinetochore heatmap

# add new cat column
dep_pe_pg_99_filt_df <- dep_pe_pg_99_filt_df %>%
    mutate(new_cat=ifelse(Gene.Names %in% cbf1_3, "cbf1_3",
                          ifelse(Gene.Names %in% inner_kt, "inner_kt",
                                 ifelse(Gene.Names %in% KMN, "KMN",
                                        ifelse(Gene.Names %in% dam1,"dam1",
                                               ifelse(Gene.Names %in% accessory, "accessory_kt",
                                                      ifelse(Gene.Names %in% maps, "maps_kt",
                                                             ifelse(Gene.Names %in% cpc_sac,"cpc_sac",
                                                                    "other"))))))))


# write a function to plot a heatmap

pe_kt_subgroup_heatmap<-function(subgroup,plot_title){
    
    dep_pe_pg_99_filt_df %>% 
    filter(new_cat==subgroup) %>% 
    select(name, all_of(pe_int_colnames)) %>% 
    # filter out proteins that have NA fc in all these columns
    # name in this case is actually a gene_name_site
    filter(if_any(-c(name),~!is.na(.x))) %>% 
    pivot_longer(-c(name),names_to="sample",values_to="log2int") %>% 
    ### the line below is important for plotting in the right order
    mutate(sample=factor(sample, levels=c("WT_mitoM_1","WT_mitoM_2","WT_mitoM_3",
                                          "WT_MII_1","WT_MII_2","WT_MII_3",
                                          "WT_MI_1","WT_MI_2","WT_MI_3",
                                          "WT_pro_1","WT_pro_2","WT_pro_3",
                                          "no_tag_mitoM_1","no_tag_mitoM_2","no_tag_mitoM_3",
                                          "no_tag_MII_1","no_tag_MII_2","no_tag_MII_3",
                                          "no_tag_MI_1","no_tag_MI_2","no_tag_MI_3",
                                          "no_tag_pro_1","no_tag_pro_2","no_tag_pro_3"))) %>%
    mutate(name=fct_reorder2(name,dplyr::desc(sample),dplyr::desc(log2int),.na_rm=F)) %>%
    mutate(int=2^log2int) %>%
    ggplot(aes(x=sample,y=name,fill=log10(int)))+
    geom_tile(color="black")+
    #scale_fill_distiller(palette="PuOr")+
    scale_fill_gradient(low="white",high="darkgreen")+
    #geom_text(aes(),size=3)+
    theme_minimal()+
    theme(axis.text.x = element_text(angle=90,vjust=0.5))+
    labs(x="",y="",fill="log10int",title=plot_title) +
    coord_flip()
}

pe_kt_cbf3_all_samples_heatmap<-pe_kt_subgroup_heatmap("cbf1_3","Cbf3 protein sites")

pe_kt_innerkt_all_samples_heatmap<-pe_kt_subgroup_heatmap("inner_kt","CCAN protein sites")

pe_kt_kmn_all_samples_heatmap<-pe_kt_subgroup_heatmap("KMN","KMN protein sites")

pe_kt_dam1_all_samples_heatmap<-pe_kt_subgroup_heatmap("dam1","Dam1c protein sites")

pe_kt_accessory_all_samples_heatmap<-pe_kt_subgroup_heatmap("accessory_kt","Accessory protein sites")

pe_kt_cpc_sac_all_samples_heatmap<-pe_kt_subgroup_heatmap("cpc_sac","CPC/SAC protein sites")

pe_kt_maps_all_samples_heatmap<-pe_kt_subgroup_heatmap("maps_kt","MAPs protein sites")


# print plots to pdf

print_plot(pe_kt_cbf3_all_samples_heatmap,"pe_kt_cbf3_all_samples_heatmap.pdf",4,4)

print_plot(pe_kt_innerkt_all_samples_heatmap,"pe_kt_innerkt_all_samples_heatmap.pdf",8,4)

print_plot(pe_kt_kmn_all_samples_heatmap,"pe_kt_kmn_all_samples_heatmap.pdf",12,4)

print_plot(pe_kt_dam1_all_samples_heatmap,"pe_kt_dam1_all_samples_heatmap.pdf",5,4)

print_plot(pe_kt_accessory_all_samples_heatmap,"pe_kt_acc_all_samples_heatmap.pdf",8,4)

print_plot(pe_kt_cpc_sac_all_samples_heatmap,"pe_kt_cpc_sac_all_samples_heatmap.pdf",8.5,4)

print_plot(pe_kt_maps_all_samples_heatmap,"pe_kt_maps_all_samples_heatmap.pdf",4.5,4)


# instead of showing replicates separately,
# plot all replicates in one boxplot,
# maybe I can color them differently for the different replicates
# add text label of number of sites in each group
n_fun <- function(x){
    return(data.frame(y = median(x,na.rm=T), label = paste0("n = ",length(!is.na(x)))))
}

# Function to plot boxplots of subcomplexes 

pe_kt_subgroup_boxplot<-function(subgroup,plot_title){
    dep_pe_pg_99_filt_df %>% 
        # filter to one sub_complex at a time otherwise the plot is too large
        filter(new_cat==subgroup) %>%
        # select only tag columns, all replicates separately
        select(name, new_cat, all_of(grep("WT",pe_int_colnames,value=T))) %>% 
        # filter out proteins that have NA fc in all these columns
        # name in this case is actually a gene_name_site
        filter(if_any(-c(name,new_cat),~!is.na(.x))) %>%
        pivot_longer(-c(name,new_cat),names_to="sample",values_to="log2int") %>% 
        mutate(stage=ifelse(grepl("mitoM",sample),"mitoM",
                            ifelse(grepl("_MII_",sample),"MII",
                                   ifelse(grepl("_MI_",sample),"MI","pro")))) %>%
        mutate(stage=factor(stage,levels=c("pro","MI","MII","mitoM"))) %>%
        mutate(replicate=ifelse(grepl("_1",sample),"rep1",
                                ifelse(grepl("_2",sample),"rep2","rep3"))) %>%
        mutate(int=2^log2int) %>%
        group_by(name, stage) %>%  # Group by gene.name.site and stage
        summarize(median_int = median(int, na.rm = TRUE), .groups = "drop") %>%
        ggplot(aes(x=stage,y=log10(median_int),fill=stage))+
        geom_boxplot(alpha=0.5)+
        stat_summary(fun.data=n_fun,geom="text")+
        #stat_summary(fun.data=n_fun,geom="boxplot",alpha=0.9)+
        #geom_jitter(aes(color=replicate), size=1, alpha=0.9) +
        theme_classic()+
        theme(axis.text.x = element_text(angle=90,vjust=0.6),
              legend.position="none")+
        labs(title=plot_title)
}

pe_kt_cbf3_boxplot<-pe_kt_subgroup_boxplot("cbf1_3","Cbf3 phosphorylation")
pe_kt_innerkt_boxplot<-pe_kt_subgroup_boxplot("inner_kt","CCAN phosphorylation")
pe_kt_kmn_boxplot<-pe_kt_subgroup_boxplot("KMN","KMN phosphorylation")
pe_kt_dam1_boxplot<-pe_kt_subgroup_boxplot("dam1","Dam1c phosphorylation")
pe_kt_acc_boxplot<-pe_kt_subgroup_boxplot("accessory_kt","Accessory KT phosphorylation")
pe_kt_cpc_sac_boxplot<-pe_kt_subgroup_boxplot("cpc_sac","CPC/SAC phosphorylation")
pe_kt_maps_boxplot<-pe_kt_subgroup_boxplot("maps_kt","MAPs phosphorylation")

# all phosphorylation boxplot

pe_total_phos_boxplot<-dep_pe_pg_99_filt_df %>% 
    # filter to one sub_complex at a time otherwise the plot is too large
    #filter(new_cat==subgroup) %>%
    # select only tag columns, all replicates separately
    select(name, all_of(grep("WT",pe_int_colnames,value=T))) %>% 
    # filter out proteins that have NA fc in all these columns
    # name in this case is actually a gene_name_site
    filter(if_any(-c(name),~!is.na(.x))) %>%
    pivot_longer(-c(name),names_to="sample",values_to="log2int") %>% 
    mutate(stage=ifelse(grepl("mitoM",sample),"mitoM",
                        ifelse(grepl("_MII_",sample),"MII",
                               ifelse(grepl("_MI_",sample),"MI","pro")))) %>%
    mutate(stage=factor(stage,levels=c("pro","MI","MII","mitoM"))) %>%
    mutate(replicate=ifelse(grepl("_1",sample),"rep1",
                            ifelse(grepl("_2",sample),"rep2","rep3"))) %>%
    mutate(int=2^log2int) %>%
    group_by(name, stage) %>%  # Group by gene.name.site and stage
    summarize(median_int = median(int, na.rm = TRUE), .groups = "drop") %>%
    ggplot(aes(x=stage,y=log10(median_int),fill=stage))+
    geom_boxplot(alpha=0.5)+
    stat_summary(fun.data=n_fun,geom="text")+
    #geom_jitter(aes(color=replicate), size=1, alpha=0.9) +
    theme_classic()+
    theme(axis.text.x = element_text(angle=90,vjust=0.6),
          legend.position="none")+
    labs(title="all phospho abundance")

#print_plot(pe_total_phos_boxplot,"pe_total_phos_boxplot.pdf",4,5)

library(ggpubr)

# takes a long time to run this
kt_boxplots<-ggarrange(pe_total_phos_boxplot,pe_kt_cbf3_boxplot,pe_kt_innerkt_boxplot,pe_kt_kmn_boxplot,
          pe_kt_dam1_boxplot,pe_kt_acc_boxplot,pe_kt_cpc_sac_boxplot,
          pe_kt_maps_boxplot,nrow=2,ncol=4,align="h")

print_plot(kt_boxplots,"kt_boxplots.pdf",12,8)

## NEW PLOT in which the control is KT protein phos instead of total phos
# create col counting whether protein is in the kinetochore list

dep_pe_pg_99_filt_df<-dep_pe_pg_99_filt_df %>%
    # create col counting whether site is in the kinetochore list
    mutate(kinetochore=ifelse(Gene.Names %in% master_kt_df$Gene,T,F))

kt_sites_boxplot<-dep_pe_pg_99_filt_df %>% 
    filter(kinetochore==TRUE) %>%
    # select only tag columns, all replicates separately
    select(name, Sequence, all_of(grep("WT",pe_int_colnames,value=T))) %>% 
    # filter out proteins that have NA fc in all these columns
    # name in this case is actually a gene_name_site
    filter(if_any(-c(name,Sequence),~!is.na(.x))) %>%
    pivot_longer(-c(name,Sequence),names_to="sample",values_to="log2int") %>% 
    mutate(stage=ifelse(grepl("mitoM",sample),"mitoM",
                        ifelse(grepl("_MII_",sample),"MII",
                               ifelse(grepl("_MI_",sample),"MI","pro")))) %>%
    mutate(stage=factor(stage,levels=c("pro","MI","MII","mitoM"))) %>%
    mutate(replicate=ifelse(grepl("_1",sample),"rep1",
                            ifelse(grepl("_2",sample),"rep2","rep3"))) %>%
    mutate(int=2^log2int) %>%
    # mutate(motif=grepl(".....[DEN].[ST].......",Sequence)) %>%
    # filter(motif==TRUE) %>%
    group_by(name, stage) %>%  # Group by gene.name.site and stage
    summarize(median_int = median(int, na.rm = TRUE), .groups = "drop") %>%
    ggplot(aes(x=stage,y=log10(median_int),fill=stage))+
    geom_boxplot(alpha=0.5)+
    stat_summary(fun.data=n_fun,geom="text")+
    stat_pwc(method="wilcox_test",label="p.adj.format",p.adjust.method = "bonferroni")+
    #geom_jitter(aes(color=replicate), size=1, alpha=0.9) +
    theme_classic()+
    theme(axis.text.x = element_text(angle=90,vjust=0.6),
          legend.position="none")+
    labs(title="kt sites")

kt_boxplots2<-ggarrange(kt_sites_boxplot,pe_kt_cbf3_boxplot,pe_kt_innerkt_boxplot,pe_kt_kmn_boxplot,
                       pe_kt_dam1_boxplot,pe_kt_acc_boxplot,pe_kt_cpc_sac_boxplot,
                       pe_kt_maps_boxplot,nrow=2,ncol=4,align="h")

print_plot(kt_boxplots2,"kt_boxplots2.pdf",12,8)


# KT subgroup boxplots with stats

stages<-c("pro","MI","MII","mitoM")
new_cat_stage<-paste0("cbf1_3.",stages)

combn(new_cat_stage,m=2,simplify=F)[[1]]

generate_list <- function(prefix) {
    pairs <- list(
        c("pro", "MI"),
        c("MI", "MII"),
        c("MII", "mitoM"),
        c("pro", "MII"),
        c("MI", "mitoM"),
        c("pro", "mitoM")
    )
    
    lapply(pairs, function(x) paste(prefix, x, sep = "."))
}

# Example usage:
generate_list("cbf1_3")
generate_list("kmn")

# kt_sub group boxplots with stats!

pe_kt_subgroup_stats_boxplot<-function(subgroup,plot_title){
    
    comparison_list<-generate_list(subgroup)

    dep_pe_pg_99_filt_df %>% 
        # filter to one sub_complex at a time otherwise the plot is too large
        filter(new_cat==subgroup) %>%
        # select only tag columns, all replicates separately
        select(name, new_cat, all_of(grep("WT",pe_int_colnames,value=T))) %>% 
        # filter out proteins that have NA fc in all these columns
        # name in this case is actually a gene_name_site
        filter(if_any(-c(name,new_cat),~!is.na(.x))) %>%
        pivot_longer(-c(name,new_cat),names_to="sample",values_to="log2int") %>% 
        mutate(stage=ifelse(grepl("mitoM",sample),"mitoM",
                            ifelse(grepl("_MII_",sample),"MII",
                                   ifelse(grepl("_MI_",sample),"MI","pro")))) %>%
        mutate(stage=factor(stage,levels=c("pro","MI","MII","mitoM"))) %>%
        mutate(replicate=ifelse(grepl("_1",sample),"rep1",
                                ifelse(grepl("_2",sample),"rep2","rep3"))) %>%
        mutate(int=2^log2int) %>%
        group_by(name, stage) %>%  # Group by gene.name.site and stage
        summarize(median_int = median(int, na.rm = TRUE), .groups = "drop") %>%
        ggplot(aes(x=stage,y=log10(median_int),fill=stage))+
        geom_boxplot(alpha=0.5)+
        stat_summary(fun.data=n_fun,geom="text")+
        stat_pwc(method="wilcox_test",label="p.adj.format",p.adjust.method = "bonferroni")+
        # stat_compare_means(comparisons=comparison_list,
        #                    method="wilcox.test",aes(label=..p.adj..))+
        #geom_jitter(aes(color=replicate), size=1, alpha=0.9) +
        theme_classic()+
        theme(axis.text.x = element_text(angle=90,vjust=0.6),
              legend.position = "none")+
        labs(title=plot_title)
}

pe_kt_cbf3_stats_boxplot<-pe_kt_subgroup_stats_boxplot("cbf1_3","Cbf3 phosphorylation")
pe_kt_innerkt_stats_boxplot<-pe_kt_subgroup_stats_boxplot("inner_kt","CCAN phosphorylation")
pe_kt_kmn_stats_boxplot<-pe_kt_subgroup_stats_boxplot("KMN","KMN phosphorylation")
pe_kt_dam1_stats_boxplot<-pe_kt_subgroup_stats_boxplot("dam1","Dam1c phosphorylation")
pe_kt_acc_stats_boxplot<-pe_kt_subgroup_stats_boxplot("accessory_kt","Accessory KT phosphorylation")
pe_kt_cpc_sac_stats_boxplot<-pe_kt_subgroup_stats_boxplot("cpc_sac","CPC/SAC phosphorylation")
pe_kt_maps_stats_boxplot<-pe_kt_subgroup_stats_boxplot("maps_kt","MAPs phosphorylation")

# all phosphorylation boxplot

# histogram of distribution of all phospho-sites at each stage
pe_total_phos_histogram<-dep_pe_pg_99_filt_df %>% 
    # filter to one sub_complex at a time otherwise the plot is too large
    #filter(new_cat==subgroup) %>%
    # select only tag columns, all replicates separately
    select(name, all_of(grep("WT",pe_int_colnames,value=T))) %>% 
    # filter out proteins that have NA fc in all these columns
    # name in this case is actually a gene_name_site
    filter(if_any(-c(name),~!is.na(.x))) %>%
    pivot_longer(-c(name),names_to="sample",values_to="log2int") %>% 
    mutate(stage=ifelse(grepl("mitoM",sample),"mitoM",
                        ifelse(grepl("_MII_",sample),"MII",
                               ifelse(grepl("_MI_",sample),"MI","pro")))) %>%
    mutate(stage=factor(stage,levels=c("pro","MI","MII","mitoM"))) %>%
    mutate(replicate=ifelse(grepl("_1",sample),"rep1",
                            ifelse(grepl("_2",sample),"rep2","rep3"))) %>%
    mutate(int=2^log2int) %>%
    # group_by(name, stage) %>%  # Group by gene.name.site and stage
    # summarize(median_int = median(int, na.rm = TRUE), .groups = "drop") %>%
    ggplot(aes(x=log10(int),fill=stage))+
    geom_histogram(bins=100)+
    facet_wrap(~stage,nrow=1)+
    theme_classic()+
    theme(axis.text.x = element_text(angle=90,vjust=0.6))+
    scale_y_continuous(expand=expansion(mult=c(0,0.2)))+
    labs(title="all phospho abundance")

print_plot(pe_total_phos_histogram,"pe_total_phos_histogram.pdf",10,4)


pe_total_phos_stats_boxplot<-dep_pe_pg_99_filt_df %>% 
    # filter to one sub_complex at a time otherwise the plot is too large
    #filter(new_cat==subgroup) %>%
    # select only tag columns, all replicates separately
    select(name, all_of(grep("WT",pe_int_colnames,value=T))) %>% 
    # filter out proteins that have NA fc in all these columns
    # name in this case is actually a gene_name_site
    filter(if_any(-c(name),~!is.na(.x))) %>%
    pivot_longer(-c(name),names_to="sample",values_to="log2int") %>% 
    mutate(stage=ifelse(grepl("mitoM",sample),"mitoM",
                        ifelse(grepl("_MII_",sample),"MII",
                               ifelse(grepl("_MI_",sample),"MI","pro")))) %>%
    mutate(stage=factor(stage,levels=c("pro","MI","MII","mitoM"))) %>%
    mutate(replicate=ifelse(grepl("_1",sample),"rep1",
                            ifelse(grepl("_2",sample),"rep2","rep3"))) %>%
    mutate(int=2^log2int) %>%
    group_by(name, stage) %>%  # Group by gene.name.site and stage
    summarize(median_int = median(int, na.rm = TRUE), .groups = "drop") %>%
    ggplot(aes(x=stage,y=log10(median_int),fill=stage))+
    geom_boxplot(alpha=0.5)+
    stat_summary(fun.data=n_fun,geom="text")+
    stat_pwc(method="wilcox_test",label="p.adj.format",p.adjust.method = "bonferroni")+
    # stat_compare_means(comparisons=list(c("pro","MI"),c("MI","MII"),
    #                                     c("MII","mitoM"),c("pro","MII"),
    #                                     c("MI","mitoM"),c("pro","mitoM")),
    #                    method="wilcox.test",aes(label=..p.adj..))+
    #geom_jitter(aes(color=replicate), size=1, alpha=0.9) +
    theme_classic()+
    theme(axis.text.x = element_text(angle=90,vjust=0.6),
          legend.position="none")+
    labs(title="all phospho abundance")

# KT only sites control # this code is already above

# kt_sites_boxplot<-dep_pe_pg_99_filt_df %>% 
#     filter(kinetochore==TRUE) %>%
#     # select only tag columns, all replicates separately
#     select(name, Sequence, all_of(grep("WT",pe_int_colnames,value=T))) %>% 
#     # filter out proteins that have NA fc in all these columns
#     # name in this case is actually a gene_name_site
#     filter(if_any(-c(name,Sequence),~!is.na(.x))) %>%
#     pivot_longer(-c(name,Sequence),names_to="sample",values_to="log2int") %>% 
#     mutate(stage=ifelse(grepl("mitoM",sample),"mitoM",
#                         ifelse(grepl("_MII_",sample),"MII",
#                                ifelse(grepl("_MI_",sample),"MI","pro")))) %>%
#     mutate(stage=factor(stage,levels=c("pro","MI","MII","mitoM"))) %>%
#     mutate(replicate=ifelse(grepl("_1",sample),"rep1",
#                             ifelse(grepl("_2",sample),"rep2","rep3"))) %>%
#     mutate(int=2^log2int) %>%
#     # mutate(motif=grepl(".....[DEN].[ST].......",Sequence)) %>%
#     # filter(motif==TRUE) %>%
#     group_by(name, stage) %>%  # Group by gene.name.site and stage
#     summarize(median_int = median(int, na.rm = TRUE), .groups = "drop") %>%
#     ggplot(aes(x=stage,y=log10(median_int),fill=stage))+
#     geom_boxplot(alpha=0.5)+
#     stat_summary(fun.data=n_fun,geom="text")+
#     stat_pwc(method="wilcox_test",label="p.adj.format",p.adjust.method = "bonferroni")+
#     #geom_jitter(aes(color=replicate), size=1, alpha=0.9) +
#     theme_classic()+
#     theme(axis.text.x = element_text(angle=90,vjust=0.6),
#           legend.position="none")+
#     labs(title="kt sites")


#print_plot(pe_total_phos_boxplot,"pe_total_phos_boxplot.pdf",4,5)

# takes a long time to run this
kt_stats_boxplots<-ggarrange(kt_sites_boxplot,pe_kt_cbf3_stats_boxplot,pe_kt_innerkt_stats_boxplot,pe_kt_kmn_stats_boxplot,
                       pe_kt_dam1_stats_boxplot,pe_kt_acc_stats_boxplot,pe_kt_cpc_sac_stats_boxplot,
                       pe_kt_maps_stats_boxplot,nrow=2,ncol=4,align="h")

print_plot(kt_stats_boxplots,"kt_stats_boxplots.pdf",15,12)


# kinetochore proteins ranked by total phosphorylation, and in each stage
pe_kt_total_phos<-dep_pe_pg_99_filt_df %>% 
    # filter to one sub_complex at a time otherwise the plot is too large
    filter(kinetochore==T) %>%
    # select only tag columns, all replicates separately
    select(name, Gene.Names,all_of(grep("WT",pe_int_colnames,value=T))) %>% 
    # filter out proteins that have NA fc in all these columns
    # name in this case is actually a gene_name_site
    filter(if_any(-c(name,Gene.Names),~!is.na(.x))) %>%
    pivot_longer(-c(name,Gene.Names),names_to="sample",values_to="log2int") %>% 
    mutate(stage=ifelse(grepl("mitoM",sample),"mitoM",
                        ifelse(grepl("_MII_",sample),"MII",
                               ifelse(grepl("_MI_",sample),"MI","pro")))) %>%
    mutate(stage=factor(stage,levels=c("pro","MI","MII","mitoM"))) %>%
    mutate(replicate=ifelse(grepl("_1",sample),"rep1",
                            ifelse(grepl("_2",sample),"rep2","rep3"))) %>%
    mutate(int=2^log2int) %>% 
    group_by(Gene.Names,stage) %>%
   mutate(gene_sum=sum(int,na.rm=T),.groups="drop") %>%
    group_by(Gene.Names) %>%
    mutate(total_gene_sum=sum(gene_sum,na.rm=T),.groups="drop") %>%
    ungroup() %>%
    mutate(Gene.Names=paste0(Gene.Names," (",log10(total_gene_sum) %>% round(digits=1),")")) %>%
    mutate(Gene.Names=fct_reorder(Gene.Names,total_gene_sum)) %>%
    # mutate(Gene.Names=fct_reorder2(Gene.Names,dplyr::desc(stage),dplyr::desc(total_gene_sum),.na_rm=T)) %>%
    #mutate(Gene.Names=fct_reorder2(Gene.Names,dplyr::desc(stage),dplyr::desc(gene_sum),.na_rm=T)) %>%
    ggplot(aes(x=stage,y=Gene.Names,fill=log10(gene_sum)))+
    geom_tile(color="black")+
    scale_fill_gradient(low="white",high="darkgreen")+
    theme_minimal()+
    theme(axis.text.x = element_text(angle=90,vjust=0.5))+
    labs(x="",y="",fill="log10int",title="total phosphorylation")

print_plot(pe_kt_total_phos,"pe_kt_total_phos.pdf",3,8)


# can I calculate which kinetochore proteins undergo 
# the most dynamic phospho changes at different stages?

# use the _diff cols which calculate pairwise fold changes between stages
# which sites have the highest median _diff (FC) ?

# only select tag cols

tag_diff_cols<-grep("_diff",colnames(dep_pe_pg_99_filt_df),value=T)[c(2,3,5,8,9,10)]

# can I add how many sites for each protein to the labels of the plot??

# KT phos dynamics ####
# REVISED dynamic range heatmap that matches the 
# intensities in the green phospho heatmap

pe_kt_dynamics_boxplot<-dep_pe_pg_99_filt_df %>% 
    # filter to one sub_complex at a time otherwise the plot is too large
    filter(kinetochore==T) %>%
    #filter(Gene.Names=="CTF13") %>%
    # only select tag columns which have "WT" in their name
    select(name, Gene.Names,all_of(grep("WT",pe_int_colnames,value=T))) %>% 
    # filter out proteins that have NA fc in all these columns
    # name in this case is actually a gene_name_site
    filter(if_any(-c(name,Gene.Names),~!is.na(.x))) %>%
    # add number of sites for each Gene column
    group_by(Gene.Names) %>%
    mutate(num_sites=n()) %>% 
    pivot_longer(-c(name,Gene.Names,num_sites),names_to="sample",values_to="int") %>% 
    # anti-log2 the int columns
    mutate(int=2^int) %>% 
    mutate(stage=ifelse(grepl("mitoM",sample),"mitoM",
                        ifelse(grepl("_MII_",sample),"MII",
                               ifelse(grepl("_MI_",sample),"MI","pro")))) %>%
    group_by(name,stage) %>%
    mutate(mean_int_stage=log10(mean(int,na.rm=T))) %>%
    # must keep "num_sites" here in order to keep the column
    #  # groups by individual sites 
    # for each Gene, what is the max range of each site across 4 stages
    # for example for 1 site it will take the max value in pro and the min in MII for ex
    group_by(name,Gene.Names,num_sites) %>%
    summarize(site_range=max(mean_int_stage,na.rm=T)-min(mean_int_stage,na.rm=T),.groups="drop") %>%
   # because some sites were only found in one stage, the range comes out to zero
   # since mean-mean = 0
   # filter out these sites
    #filter(site_range>0) %>% 
    mutate(Gene.Names=paste0(Gene.Names," (",num_sites,")")) %>%
    mutate(Gene.Names=fct_reorder(Gene.Names,site_range,na.rm=T)) %>%
    ggplot(aes(x=Gene.Names,y=(site_range)))+
    geom_boxplot()+
    geom_jitter(alpha=0.5)+
    coord_flip()+
    labs(x="",y="max site range")+
    theme_classic()+
    theme(axis.text.y=element_text(size=rel(1.5)))

print_plot(pe_kt_dynamics_boxplot,"pe_kt_dynamics_boxplot.pdf",4,9)

# boxplots of individual sites that seem interesting

sel_site_bars<-dep_pe_pg_99_filt_df %>% 
    filter(name %in% c("DSN1_69","OKP1_70","NDC80_54","SLK19_23")) %>%
    # add residue to the name col
    mutate(name=paste0(Gene.Names,"_",Residue,"_",Site)) %>%
    # select only tag columns, all replicates separately
    select(name, Site, all_of(grep("WT",pe_int_colnames,value=T))) %>% 
    # filter out proteins that have NA fc in all these columns
    # name in this case is actually a gene_name_site
    filter(if_any(-c(name,Site),~!is.na(.x))) %>%
    pivot_longer(-c(name,Site),names_to="sample",values_to="log2int") %>% 
    mutate(stage=ifelse(grepl("mitoM",sample),"mitoM",
                        ifelse(grepl("_MII_",sample),"MII",
                               ifelse(grepl("_MI_",sample),"MI","pro")))) %>%
    mutate(stage=factor(stage,levels=c("pro","MI","MII","mitoM"))) %>%
    mutate(replicate=ifelse(grepl("_1",sample),"rep1",
                            ifelse(grepl("_2",sample),"rep2","rep3"))) %>%
    #mutate(int=2^log2int) %>% 
    mutate(name=fct_reorder(name,Site,.fun=min)) %>%
    ggplot(aes(x=stage,y=log2int,fill=stage))+
    geom_bar(stat="summary", fun="mean", position="dodge",alpha=0.8) +  # Use mean log2int for bars
    geom_point(aes(color=replicate), size=2, position=position_dodge(width=0.9)) +  # Align points
    facet_wrap(~name)+
    #stat_summary(fun.data=n_fun,geom="text")+
    stat_pwc(method="t_test",label="p.adj.format",p.adjust.method = "bonferroni")+
    theme_classic()+
    theme(axis.text.x = element_text(angle=90,vjust=0.6))+
    labs(title="selected phospho-sites")

print_plot(sel_site_bars,"sel_site_bars.pdf",5,8) 

# REVISED phospho-motif analysis to match ####
# KT categories from preivous figure

library(ggseqlogo)

pe_col_names<-colnames(dep_pe_pg_99_filt_df[,2:23])

MII_tag_cols<-pe_col_names[1:3]
MII_notag_cols<-pe_col_names[4:6]
pro_tag_cols<-pe_col_names[7:9]
pro_notag_cols<-pe_col_names[10:12]
MI_tag_cols<-pe_col_names[13:15]
MI_notag_cols<-pe_col_names[16:18]
mitoM_tag_cols<-pe_col_names[19:20]
mitoM_notag_cols<-pe_col_names[21:22]

new_pro_logo<-dep_pe_pg_99_filt_df %>%
    filter(kinetochore==TRUE) %>%
    # select only tag columns, all replicates separately
    select(name, Sequence,all_of(grep("WT",pe_int_colnames,value=T))) %>% 
    # filter out proteins that have NA in all these columns
    filter(if_any(-name,~!is.na(.x))) %>%
    pivot_longer(-c(name,Sequence),names_to="sample",values_to="log2int") %>%
    mutate(stage=ifelse(grepl("mitoM",sample),"mitoM",
                        ifelse(grepl("_MII_",sample),"MII",
                               ifelse(grepl("_MI_",sample),"MI","pro")))) %>%
    mutate(stage=factor(stage,levels=c("pro","MI","MII","mitoM"))) %>%
    mutate(replicate=ifelse(grepl("_1",sample),"rep1",
                            ifelse(grepl("_2",sample),"rep2","rep3"))) %>%
    mutate(int=2^log2int) %>% 
    group_by(name, stage, Sequence) %>%  # Group by gene.name.site and stage
    summarize(median_int = median(int, na.rm = TRUE), .groups = "drop") %>%
    filter(stage=="pro") %>%
    filter(!is.na(median_int)) %>%
    distinct(Sequence) %>% #nrow() #100 as expected
    unlist() %>%
    substr(.,3,13) %>%
    ggseqlogo(.,method="prob")+
    ggtitle("prophase kt sites, n=100")+
    theme(legend.position="none")+
    scale_x_continuous(breaks=1:11,labels=-5:5)

# MI
new_MI_logo<-dep_pe_pg_99_filt_df %>%
    filter(kinetochore==TRUE) %>%
    # select only tag columns, all replicates separately
    select(name, Sequence,all_of(grep("WT",pe_int_colnames,value=T))) %>% 
    # filter out proteins that have NA in all these columns
    filter(if_any(-name,~!is.na(.x))) %>%
    pivot_longer(-c(name,Sequence),names_to="sample",values_to="log2int") %>%
    mutate(stage=ifelse(grepl("mitoM",sample),"mitoM",
                        ifelse(grepl("_MII_",sample),"MII",
                               ifelse(grepl("_MI_",sample),"MI","pro")))) %>%
    mutate(stage=factor(stage,levels=c("pro","MI","MII","mitoM"))) %>%
    mutate(replicate=ifelse(grepl("_1",sample),"rep1",
                            ifelse(grepl("_2",sample),"rep2","rep3"))) %>%
    mutate(int=2^log2int) %>% 
    group_by(name, stage, Sequence) %>%  # Group by gene.name.site and stage
    summarize(median_int = median(int, na.rm = TRUE), .groups = "drop") %>%
    filter(stage=="MI") %>%
    filter(!is.na(median_int)) %>%
    distinct(Sequence) %>% #nrow() #100 as expected
    unlist() %>%
    substr(.,3,13) %>%
    ggseqlogo(.,method="prob")+
    ggtitle("MI kt sites, n=143")+
    theme(legend.position="none")+
    scale_x_continuous(breaks=1:11,labels=-5:5)

# MII
new_MII_logo<-dep_pe_pg_99_filt_df %>%
    filter(kinetochore==TRUE) %>%
    # select only tag columns, all replicates separately
    select(name, Sequence,all_of(grep("WT",pe_int_colnames,value=T))) %>% 
    # filter out proteins that have NA in all these columns
    filter(if_any(-name,~!is.na(.x))) %>%
    pivot_longer(-c(name,Sequence),names_to="sample",values_to="log2int") %>%
    mutate(stage=ifelse(grepl("mitoM",sample),"mitoM",
                        ifelse(grepl("_MII_",sample),"MII",
                               ifelse(grepl("_MI_",sample),"MI","pro")))) %>%
    mutate(stage=factor(stage,levels=c("pro","MI","MII","mitoM"))) %>%
    mutate(replicate=ifelse(grepl("_1",sample),"rep1",
                            ifelse(grepl("_2",sample),"rep2","rep3"))) %>%
    mutate(int=2^log2int) %>% 
    group_by(name, stage, Sequence) %>%  # Group by gene.name.site and stage
    summarize(median_int = median(int, na.rm = TRUE), .groups = "drop") %>%
    filter(stage=="MII") %>%
    filter(!is.na(median_int)) %>%
    distinct(Sequence) %>% #nrow() #100 as expected
    unlist() %>%
    substr(.,3,13) %>%
    ggseqlogo(.,method="prob")+
    ggtitle("MII kt sites, n=119")+
    theme(legend.position="none")+
    scale_x_continuous(breaks=1:11,labels=-5:5)

# mitoM
new_mitoM_logo<-dep_pe_pg_99_filt_df %>%
    filter(kinetochore==TRUE) %>%
    # select only tag columns, all replicates separately
    select(name, Sequence,all_of(grep("WT",pe_int_colnames,value=T))) %>% 
    # filter out proteins that have NA in all these columns
    filter(if_any(-name,~!is.na(.x))) %>%
    pivot_longer(-c(name,Sequence),names_to="sample",values_to="log2int") %>%
    mutate(stage=ifelse(grepl("mitoM",sample),"mitoM",
                        ifelse(grepl("_MII_",sample),"MII",
                               ifelse(grepl("_MI_",sample),"MI","pro")))) %>%
    mutate(stage=factor(stage,levels=c("pro","MI","MII","mitoM"))) %>%
    mutate(replicate=ifelse(grepl("_1",sample),"rep1",
                            ifelse(grepl("_2",sample),"rep2","rep3"))) %>%
    mutate(int=2^log2int) %>% 
    group_by(name, stage, Sequence) %>%  # Group by gene.name.site and stage
    summarize(median_int = median(int, na.rm = TRUE), .groups = "drop") %>%
    filter(stage=="mitoM") %>%
    filter(!is.na(median_int)) %>%
    distinct(Sequence) %>% #nrow() #100 as expected
    unlist() %>%
    substr(.,3,13) %>%
    ggseqlogo(.,method="prob")+
    ggtitle("mitoM kt sites, n=124")+
    theme(legend.position="none")+
    scale_x_continuous(breaks=1:11,labels=-5:5)

new_detected_kt_site_logos<-ggarrange(new_pro_logo,new_MI_logo,
                                      new_MII_logo,new_mitoM_logo)

print_plot(new_detected_kt_site_logos,"new_detected_kt_site_logos.pdf",8,6)
    
# group_by(stage) %>%
#     summarize(n=sum(!is.na(median_int))) #100,143,119,124 same numbers as boxplots

# REVISED motif counts ####

detected_df<-dep_pe_pg_99_filt_df %>%
    filter(kinetochore==TRUE) %>%
    # select only tag columns, all replicates separately
    select(name, Sequence,all_of(grep("WT",pe_int_colnames,value=T))) %>% 
    # filter out proteins that have NA in all these columns
    filter(if_any(-name,~!is.na(.x))) %>%
    pivot_longer(-c(name,Sequence),names_to="sample",values_to="log2int") %>%
    mutate(stage=ifelse(grepl("mitoM",sample),"mitoM",
                        ifelse(grepl("_MII_",sample),"MII",
                               ifelse(grepl("_MI_",sample),"MI","pro")))) %>%
    mutate(stage=factor(stage,levels=c("pro","MI","MII","mitoM"))) %>%
    mutate(replicate=ifelse(grepl("_1",sample),"rep1",
                            ifelse(grepl("_2",sample),"rep2","rep3"))) %>%
    mutate(int=2^log2int) %>% 
    group_by(name, stage, Sequence) %>%  # Group by gene.name.site and stage
    summarize(median_int = median(int, na.rm = TRUE), .groups = "drop") %>%
    group_by(stage) %>%
    mutate(detected=!is.na(median_int)) %>% 
    # head()
    # group_by(stage) %>%
    #     summarize(n=sum(!is.na(median_int))) #100,143,119,124 same numbers as boxplots
    select(stage,Sequence,detected) 


new_SP_kt_sites_bar<-detected_df %>%
    filter(detected==TRUE) %>%
    # create col that says whether seq matches motif or not
    mutate(motif=grepl(".......[ST]P......",Sequence)) %>% 
    group_by(stage) %>%
    summarise(
        total_sequences = n(),
        motif_sequences = sum(motif)
    )  %>%
    mutate(perc_match=motif_sequences/total_sequences*100) %>%
    mutate(stage=factor(stage,levels=c("pro","MI","MII","mitoM"))) %>%
    #select(stage,perc_match) %>% head(10)
    ggplot(aes(x=stage,y=perc_match))+
    geom_bar(stat="identity")+
    scale_y_continuous(expand=expansion(mult=c(0,0.2)))+ # 20% more plot area after max value
    #stat_pwc(method="wilcox_test",label="p.adj.format",p.adjust.method = "bonferroni")+
    theme_minimal()+
    theme(axis.text.x = element_text(angle=90,vjust=0.5),legend.position="none")+
    labs(x="",y="% match",title="kt [ST]*P")

new_SPxKR_kt_sites_bar<-detected_df %>%
    filter(detected==TRUE) %>%
    # create col that says whether seq matches motif or not
    mutate(motif=grepl(".......[ST]P.[KR]....",Sequence)) %>% 
    group_by(stage) %>%
    summarise(
        total_sequences = n(),
        motif_sequences = sum(motif)
    )  %>%
    mutate(perc_match=motif_sequences/total_sequences*100) %>%
    mutate(stage=factor(stage,levels=c("pro","MI","MII","mitoM"))) %>%
    #select(stage,perc_match) %>% head(10)
    ggplot(aes(x=stage,y=perc_match))+
    geom_bar(stat="identity")+
    scale_y_continuous(expand=expansion(mult=c(0,0.2)))+ # 20% more plot area after max value
    #stat_pwc(method="wilcox_test",label="p.adj.format",p.adjust.method = "bonferroni")+
    theme_minimal()+
    theme(axis.text.x = element_text(angle=90,vjust=0.5),legend.position="none")+
    labs(x="",y="% match",title="kt [ST]*Px[KR]")

new_RKxST_kt_sites_bar<-detected_df %>%
    filter(detected==TRUE) %>%
    # create col that says whether seq matches motif or not
    mutate(motif=grepl(".....[RK].[ST].......",Sequence)) %>% 
    group_by(stage) %>%
    summarise(
        total_sequences = n(),
        motif_sequences = sum(motif)
    )  %>%
    mutate(perc_match=motif_sequences/total_sequences*100) %>%
    mutate(stage=factor(stage,levels=c("pro","MI","MII","mitoM"))) %>%
    #select(stage,perc_match) %>% head(10)
    ggplot(aes(x=stage,y=perc_match))+
    geom_bar(stat="identity")+
    scale_y_continuous(expand=expansion(mult=c(0,0.2)))+ # 20% more plot area after max value
    #stat_pwc(method="wilcox_test",label="p.adj.format",p.adjust.method = "bonferroni")+
    theme_minimal()+
    theme(axis.text.x = element_text(angle=90,vjust=0.5),legend.position="none")+
    labs(x="",y="% match",title="kt [RK]x[ST]*")

new_DENxST_kt_sites_bar<-detected_df %>%
    filter(detected==TRUE) %>%
    # create col that says whether seq matches motif or not
    mutate(motif=grepl(".....[DEN].[ST].......",Sequence)) %>% 
    group_by(stage) %>%
    summarise(
        total_sequences = n(),
        motif_sequences = sum(motif)
    )  %>%
    mutate(perc_match=motif_sequences/total_sequences*100) %>%
    mutate(stage=factor(stage,levels=c("pro","MI","MII","mitoM"))) %>%
    #select(stage,perc_match) %>% head(10)
    ggplot(aes(x=stage,y=perc_match))+
    geom_bar(stat="identity")+
    scale_y_continuous(expand=expansion(mult=c(0,0.2)))+ # 20% more plot area after max value
    #stat_pwc(method="wilcox_test",label="p.adj.format",p.adjust.method = "bonferroni")+
    theme_minimal()+
    theme(axis.text.x = element_text(angle=90,vjust=0.5),legend.position="none")+
    labs(x="",y="% match",title="kt [DEN]x[ST]*")

new_DENxSTF_kt_sites_bar<-detected_df %>%
    filter(detected==TRUE) %>%
    # create col that says whether seq matches motif or not
    mutate(motif=grepl(".....[DEN].[ST]F......",Sequence)) %>% 
    group_by(stage) %>%
    summarise(
        total_sequences = n(),
        motif_sequences = sum(motif)
    )  %>%
    mutate(perc_match=motif_sequences/total_sequences*100) %>%
    mutate(stage=factor(stage,levels=c("pro","MI","MII","mitoM"))) %>%
    #select(stage,perc_match) %>% head(10)
    ggplot(aes(x=stage,y=perc_match))+
    geom_bar(stat="identity")+
    scale_y_continuous(expand=expansion(mult=c(0,0.2)))+ # 20% more plot area after max value
    #stat_pwc(method="wilcox_test",label="p.adj.format",p.adjust.method = "bonferroni")+
    theme_minimal()+
    theme(axis.text.x = element_text(angle=90,vjust=0.5),legend.position="none")+
    labs(x="",y="% match",title="kt [DEN]x[ST]*F")

new_STTSDE_kt_sites_bar<-detected_df %>%
    filter(detected==TRUE) %>%
    # create col that says whether seq matches motif or not
    mutate(motif=grepl(".......[ST][TSDE]......",Sequence)) %>% 
    group_by(stage) %>%
    summarise(
        total_sequences = n(),
        motif_sequences = sum(motif)
    )  %>%
    mutate(perc_match=motif_sequences/total_sequences*100) %>%
    mutate(stage=factor(stage,levels=c("pro","MI","MII","mitoM"))) %>%
    #select(stage,perc_match) %>% head(10)
    ggplot(aes(x=stage,y=perc_match))+
    geom_bar(stat="identity")+
    scale_y_continuous(expand=expansion(mult=c(0,0.2)))+ # 20% more plot area after max value
    #stat_pwc(method="wilcox_test",label="p.adj.format",p.adjust.method = "bonferroni")+
    theme_minimal()+
    theme(axis.text.x = element_text(angle=90,vjust=0.5),legend.position="none")+
    labs(x="",y="% match",title="kt [ST]*[STDE]")

# new plot with 2x more motifs
print_plot(ggarrange(new_SP_kt_sites_bar,new_SPxKR_kt_sites_bar,new_RKxST_kt_sites_bar,new_DENxST_kt_sites_bar,new_DENxSTF_kt_sites_bar,
                     new_STTSDE_kt_sites_bar, nrow=2,ncol=3),"new_kt_sites_bar2.pdf",7,5)

# core vs accessory KT phospho-sites

dep_pe_pg_99_filt_df %>%
    group_by(new_cat) %>%
    summarize(n=n())


mitoM_core_kt_sites_logo<-dep_pe_pg_99_filt_df %>%
    filter(new_cat=="KMN"|
               new_cat=="cbf1_3"|
               new_cat=="dam1"|
               new_cat=="inner_kt") %>%
    filter(detect_mitoM) %>%
    select(Sequence) %>%
    unlist() %>%
    substr(.,3,13) %>%
    ggseqlogo(.,method="prob")+
    ggtitle(paste0("mitoM core KT sites, n=",nrow(dep_pe_pg_99_filt_df %>%
                                                      filter(new_cat=="KMN"|
                                                                 new_cat=="cbf1_3"|
                                                                 new_cat=="dam1"|
                                                                 new_cat=="inner_kt") %>% 
                                              filter(detect_mitoM))))+
    theme(legend.position="none")+
    scale_x_continuous(breaks=1:11,labels=-5:5)

pro_core_kt_sites_logo<-dep_pe_pg_99_filt_df %>%
    filter(new_cat=="KMN"|
               new_cat=="cbf1_3"|
               new_cat=="dam1"|
               new_cat=="inner_kt") %>%
    filter(detect_pro) %>%
    select(Sequence) %>%
    unlist() %>%
    substr(.,3,13) %>%
    ggseqlogo(.,method="prob")+
    ggtitle(paste0("pro core KT sites, n=",nrow(dep_pe_pg_99_filt_df %>%
                                                      filter(new_cat=="KMN"|
                                                                 new_cat=="cbf1_3"|
                                                                 new_cat=="dam1"|
                                                                 new_cat=="inner_kt") %>% 
                                                      filter(detect_pro))))+
    theme(legend.position="none")+
    scale_x_continuous(breaks=1:11,labels=-5:5)

MI_core_kt_sites_logo<-dep_pe_pg_99_filt_df %>%
    filter(new_cat=="KMN"|
               new_cat=="cbf1_3"|
               new_cat=="dam1"|
               new_cat=="inner_kt") %>%
    filter(detect_MI) %>%
    select(Sequence) %>%
    unlist() %>%
    substr(.,3,13) %>%
    ggseqlogo(.,method="prob")+
    ggtitle(paste0("MI core KT sites, n=",nrow(dep_pe_pg_99_filt_df %>%
                                                      filter(new_cat=="KMN"|
                                                                 new_cat=="cbf1_3"|
                                                                 new_cat=="dam1"|
                                                                 new_cat=="inner_kt") %>% 
                                                      filter(detect_MI))))+
    theme(legend.position="none")+
    scale_x_continuous(breaks=1:11,labels=-5:5)

MII_core_kt_sites_logo<-dep_pe_pg_99_filt_df %>%
    filter(new_cat=="KMN"|
               new_cat=="cbf1_3"|
               new_cat=="dam1"|
               new_cat=="inner_kt") %>%
    filter(detect_MII) %>%
    select(Sequence) %>%
    unlist() %>%
    substr(.,3,13) %>%
    ggseqlogo(.,method="prob")+
    ggtitle(paste0("MII core KT sites, n=",nrow(dep_pe_pg_99_filt_df %>%
                                                      filter(new_cat=="KMN"|
                                                                 new_cat=="cbf1_3"|
                                                                 new_cat=="dam1"|
                                                                 new_cat=="inner_kt") %>% 
                                                      filter(detect_MII))))+
    theme(legend.position="none")+
    scale_x_continuous(breaks=1:11,labels=-5:5)

detected_core_kt_site_logos<-ggarrange(pro_core_kt_sites_logo,MI_core_kt_sites_logo,
                                  MII_core_kt_sites_logo,mitoM_core_kt_sites_logo)

print_plot(detected_core_kt_site_logos,"detected_core_kt_site_logos.pdf",8,6)

# inner vs outer kinetochore
# subcomplex motif trends??

MI_core_kt_logo<-function(subcomplex_list,title){
    dep_pe_pg_99_filt_df %>%
        filter(new_cat %in% subcomplex_list) %>%
        filter(detect_MI) %>%
        select(Sequence) %>%
        unlist() %>%
        substr(.,3,13) %>%
        ggseqlogo(.,method="prob")+
        ggtitle(paste0(title," n=",nrow(dep_pe_pg_99_filt_df %>%
                                                   filter(new_cat %in% subcomplex_list) %>% 
                                                          filter(detect_MI))))+
        theme(legend.position="none")+
        scale_x_continuous(breaks=1:11,labels=-5:5)
}

MII_core_kt_logo<-function(subcomplex_list,title){
    dep_pe_pg_99_filt_df %>%
        filter(new_cat %in% subcomplex_list) %>%
        filter(detect_MII) %>%
        select(Sequence) %>%
        unlist() %>%
        substr(.,3,13) %>%
        ggseqlogo(.,method="prob")+
        ggtitle(paste0(title," n=",nrow(dep_pe_pg_99_filt_df %>%
                                                   filter(new_cat %in% subcomplex_list) %>% 
                                                   filter(detect_MII))))+
        theme(legend.position="none")+
        scale_x_continuous(breaks=1:11,labels=-5:5)
}

pro_core_kt_logo<-function(subcomplex_list,title){
    dep_pe_pg_99_filt_df %>%
        filter(new_cat %in% subcomplex_list) %>%
        filter(detect_pro) %>%
        select(Sequence) %>%
        unlist() %>%
        substr(.,3,13) %>%
        ggseqlogo(.,method="prob")+
        ggtitle(paste0(title," n=",nrow(dep_pe_pg_99_filt_df %>%
                                                   filter(new_cat %in% subcomplex_list) %>% 
                                                   filter(detect_pro))))+
        theme(legend.position="none")+
        scale_x_continuous(breaks=1:11,labels=-5:5)
}

mitoM_core_kt_logo<-function(subcomplex_list,title){
    dep_pe_pg_99_filt_df %>%
        filter(new_cat %in% subcomplex_list) %>%
        filter(detect_mitoM) %>%
        select(Sequence) %>%
        unlist() %>%
        substr(.,3,13) %>%
        ggseqlogo(.,method="prob")+
        ggtitle(paste0(title," n=",nrow(dep_pe_pg_99_filt_df %>%
                                                   filter(new_cat %in% subcomplex_list) %>% 
                                                   filter(detect_mitoM))))+
        theme(legend.position="none")+
        scale_x_continuous(breaks=1:11,labels=-5:5)
}

print_plot(ggarrange(
pro_core_kt_logo(c("cbf1_3","inner_kt"),"pro inner sites"),
pro_core_kt_logo(c("KMN","dam1"),"pro outer sites"),
MI_core_kt_logo(c("cbf1_3","inner_kt"),"MI inner sites"),
MI_core_kt_logo(c("KMN","dam1"),"MI outer sites"),
MII_core_kt_logo(c("cbf1_3","inner_kt"),"MII inner sites"),
MII_core_kt_logo(c("KMN","dam1"),"MII outer sites"),
mitoM_core_kt_logo(c("cbf1_3","inner_kt"),"mitoM inner sites"),
mitoM_core_kt_logo(c("KMN","dam1"),"mitoM outer sites"), nrow=4,ncol=2),
"core_kt_inner_outer_logos.pdf",10,12)

# intensity of motif sites ####

# boxplot showing intensity of motif-matching sites at each stage
# start with all sites then show KT sites

all_SP_boxplot<-dep_pe_pg_99_filt_df %>% 
    # select only tag columns, all replicates separately
    select(name, Sequence, all_of(grep("WT",pe_int_colnames,value=T))) %>% 
    # filter out proteins that have NA fc in all these columns
    # name in this case is actually a gene_name_site
    filter(if_any(-c(name,Sequence),~!is.na(.x))) %>%
    pivot_longer(-c(name,Sequence),names_to="sample",values_to="log2int") %>% 
    mutate(stage=ifelse(grepl("mitoM",sample),"mitoM",
                        ifelse(grepl("_MII_",sample),"MII",
                               ifelse(grepl("_MI_",sample),"MI","pro")))) %>%
    mutate(stage=factor(stage,levels=c("pro","MI","MII","mitoM"))) %>%
    mutate(replicate=ifelse(grepl("_1",sample),"rep1",
                            ifelse(grepl("_2",sample),"rep2","rep3"))) %>%
    mutate(int=2^log2int) %>%
    mutate(motif=grepl(".......[ST]P......",Sequence)) %>%
    filter(motif==TRUE) %>%
    group_by(name, stage) %>%  # Group by gene.name.site and stage
    summarize(median_int = median(int, na.rm = TRUE), .groups = "drop") %>%
    ggplot(aes(x=stage,y=log10(median_int),fill=stage))+
    geom_boxplot(alpha=0.5)+
    stat_summary(fun.data=n_fun,geom="text")+
    stat_pwc(method="wilcox_test",label="p.adj.format",p.adjust.method = "bonferroni")+
    #geom_jitter(aes(color=replicate), size=1, alpha=0.9) +
    theme_classic()+
    theme(axis.text.x = element_text(angle=90,vjust=0.6),
          legend.position="none")+
    labs(title="SP sites")

all_DEN_boxplot<-dep_pe_pg_99_filt_df %>% 
    # select only tag columns, all replicates separately
    select(name, Sequence, all_of(grep("WT",pe_int_colnames,value=T))) %>% 
    # filter out proteins that have NA fc in all these columns
    # name in this case is actually a gene_name_site
    filter(if_any(-c(name,Sequence),~!is.na(.x))) %>%
    pivot_longer(-c(name,Sequence),names_to="sample",values_to="log2int") %>% 
    mutate(stage=ifelse(grepl("mitoM",sample),"mitoM",
                        ifelse(grepl("_MII_",sample),"MII",
                               ifelse(grepl("_MI_",sample),"MI","pro")))) %>%
    mutate(stage=factor(stage,levels=c("pro","MI","MII","mitoM"))) %>%
    mutate(replicate=ifelse(grepl("_1",sample),"rep1",
                            ifelse(grepl("_2",sample),"rep2","rep3"))) %>%
    mutate(int=2^log2int) %>%
    mutate(motif=grepl(".....[DEN].[ST].......",Sequence)) %>%
    filter(motif==TRUE) %>%
    group_by(name, stage) %>%  # Group by gene.name.site and stage
    summarize(median_int = median(int, na.rm = TRUE), .groups = "drop") %>%
    ggplot(aes(x=stage,y=log10(median_int),fill=stage))+
    geom_boxplot(alpha=0.5)+
    stat_summary(fun.data=n_fun,geom="text")+
    stat_pwc(method="wilcox_test",label="p.adj.format",p.adjust.method = "bonferroni")+
    #geom_jitter(aes(color=replicate), size=1, alpha=0.9) +
    theme_classic()+
    theme(axis.text.x = element_text(angle=90,vjust=0.6),
          legend.position="none")+
    labs(title="DEN sites")

print_plot(ggarrange(pe_total_phos_boxplot+theme(legend.position="none")+
                         stat_pwc(method="wilcox_test",label="p.adj.format",p.adjust.method = "bonferroni"),all_SP_boxplot,
          all_DEN_boxplot,nrow=1),
          "all_motif_sites_boxplot.pdf",12,5)

# just KT phospho sites

kt_SP_boxplot<-dep_pe_pg_99_filt_df %>% 
    filter(kinetochore==T) %>%
    # select only tag columns, all replicates separately
    select(name, Sequence, all_of(grep("WT",pe_int_colnames,value=T))) %>% 
    # filter out proteins that have NA fc in all these columns
    # name in this case is actually a gene_name_site
    filter(if_any(-c(name,Sequence),~!is.na(.x))) %>%
    pivot_longer(-c(name,Sequence),names_to="sample",values_to="log2int") %>% 
    mutate(stage=ifelse(grepl("mitoM",sample),"mitoM",
                        ifelse(grepl("_MII_",sample),"MII",
                               ifelse(grepl("_MI_",sample),"MI","pro")))) %>%
    mutate(stage=factor(stage,levels=c("pro","MI","MII","mitoM"))) %>%
    mutate(replicate=ifelse(grepl("_1",sample),"rep1",
                            ifelse(grepl("_2",sample),"rep2","rep3"))) %>%
    mutate(int=2^log2int) %>%
    mutate(motif=grepl(".......[ST]P......",Sequence)) %>%
    filter(motif==TRUE) %>%
    group_by(name, stage) %>%  # Group by gene.name.site and stage
    summarize(median_int = median(int, na.rm = TRUE), .groups = "drop") %>%
    # double checking this has the correct
    # totala numbre of KT phos sites -- 100, 143,119,124
    # it does
    # group_by(stage) %>%
    # summarize(n=sum(!is.na(median_int)))
    ggplot(aes(x=stage,y=log10(median_int),fill=stage))+
    geom_boxplot(alpha=0.5)+
    stat_summary(fun.data=n_fun,geom="text")+
    stat_pwc(method="wilcox_test",label="p.adj.format",p.adjust.method = "bonferroni")+
    #geom_jitter(aes(color=replicate), size=1, alpha=0.9) +
    theme_classic()+
    theme(axis.text.x = element_text(angle=90,vjust=0.6),
          legend.position="none")+
    labs(title="kt SP sites")

kt_SPxKR_boxplot<-dep_pe_pg_99_filt_df %>% 
    filter(kinetochore==T) %>%
    # select only tag columns, all replicates separately
    select(name, Sequence, all_of(grep("WT",pe_int_colnames,value=T))) %>% 
    # filter out proteins that have NA fc in all these columns
    # name in this case is actually a gene_name_site
    filter(if_any(-c(name,Sequence),~!is.na(.x))) %>%
    pivot_longer(-c(name,Sequence),names_to="sample",values_to="log2int") %>% 
    mutate(stage=ifelse(grepl("mitoM",sample),"mitoM",
                        ifelse(grepl("_MII_",sample),"MII",
                               ifelse(grepl("_MI_",sample),"MI","pro")))) %>%
    mutate(stage=factor(stage,levels=c("pro","MI","MII","mitoM"))) %>%
    mutate(replicate=ifelse(grepl("_1",sample),"rep1",
                            ifelse(grepl("_2",sample),"rep2","rep3"))) %>%
    mutate(int=2^log2int) %>%
    mutate(motif=grepl(".......[ST]P.[KR]....",Sequence)) %>%
    filter(motif==TRUE) %>%
    group_by(name, stage) %>%  # Group by gene.name.site and stage
    summarize(median_int = median(int, na.rm = TRUE), .groups = "drop") %>%
    # double checking this has the correct
    # totala numbre of KT phos sites -- 100, 143,119,124
    # it does
    # group_by(stage) %>%
    # summarize(n=sum(!is.na(median_int)))
    ggplot(aes(x=stage,y=log10(median_int),fill=stage))+
    geom_boxplot(alpha=0.5)+
    stat_summary(fun.data=n_fun,geom="text")+
    stat_pwc(method="wilcox_test",label="p.adj.format",p.adjust.method = "bonferroni")+
    #geom_jitter(aes(color=replicate), size=1, alpha=0.9) +
    theme_classic()+
    theme(axis.text.x = element_text(angle=90,vjust=0.6),
          legend.position="none")+
    labs(title="kt SPxKR sites")

kt_DEN_boxplot<-dep_pe_pg_99_filt_df %>% 
    filter(kinetochore==TRUE) %>%
    # select only tag columns, all replicates separately
    select(name, Sequence, all_of(grep("WT",pe_int_colnames,value=T))) %>% 
    # filter out proteins that have NA fc in all these columns
    # name in this case is actually a gene_name_site
    filter(if_any(-c(name,Sequence),~!is.na(.x))) %>%
    pivot_longer(-c(name,Sequence),names_to="sample",values_to="log2int") %>% 
    mutate(stage=ifelse(grepl("mitoM",sample),"mitoM",
                        ifelse(grepl("_MII_",sample),"MII",
                               ifelse(grepl("_MI_",sample),"MI","pro")))) %>%
    mutate(stage=factor(stage,levels=c("pro","MI","MII","mitoM"))) %>%
    mutate(replicate=ifelse(grepl("_1",sample),"rep1",
                            ifelse(grepl("_2",sample),"rep2","rep3"))) %>%
    mutate(int=2^log2int) %>%
    mutate(motif=grepl(".....[DEN].[ST].......",Sequence)) %>%
    filter(motif==TRUE) %>%
    group_by(name, stage) %>%  # Group by gene.name.site and stage
    summarize(median_int = median(int, na.rm = TRUE), .groups = "drop") %>%
    ggplot(aes(x=stage,y=log10(median_int),fill=stage))+
    geom_boxplot(alpha=0.5)+
    stat_summary(fun.data=n_fun,geom="text")+
    stat_pwc(method="wilcox_test",label="p.adj.format",p.adjust.method = "bonferroni")+
    #geom_jitter(aes(color=replicate), size=1, alpha=0.9) +
    theme_classic()+
    theme(axis.text.x = element_text(angle=90,vjust=0.6),
          legend.position="none")+
    labs(title="kt DEN sites")

kt_DENF_boxplot<-dep_pe_pg_99_filt_df %>% 
    filter(kinetochore==TRUE) %>%
    # select only tag columns, all replicates separately
    select(name, Sequence, all_of(grep("WT",pe_int_colnames,value=T))) %>% 
    # filter out proteins that have NA fc in all these columns
    # name in this case is actually a gene_name_site
    filter(if_any(-c(name,Sequence),~!is.na(.x))) %>%
    pivot_longer(-c(name,Sequence),names_to="sample",values_to="log2int") %>% 
    mutate(stage=ifelse(grepl("mitoM",sample),"mitoM",
                        ifelse(grepl("_MII_",sample),"MII",
                               ifelse(grepl("_MI_",sample),"MI","pro")))) %>%
    mutate(stage=factor(stage,levels=c("pro","MI","MII","mitoM"))) %>%
    mutate(replicate=ifelse(grepl("_1",sample),"rep1",
                            ifelse(grepl("_2",sample),"rep2","rep3"))) %>%
    mutate(int=2^log2int) %>%
    mutate(motif=grepl(".....[DEN].[ST]F......",Sequence)) %>%
    filter(motif==TRUE) %>%
    group_by(name, stage) %>%  # Group by gene.name.site and stage
    summarize(median_int = median(int, na.rm = TRUE), .groups = "drop") %>%
    ggplot(aes(x=stage,y=log10(median_int),fill=stage))+
    geom_boxplot(alpha=0.5)+
    stat_summary(fun.data=n_fun,geom="text")+
    stat_pwc(method="wilcox_test",label="p.adj.format",p.adjust.method = "bonferroni")+
    #geom_jitter(aes(color=replicate), size=1, alpha=0.9) +
    theme_classic()+
    theme(axis.text.x = element_text(angle=90,vjust=0.6),
          legend.position="none")+
    labs(title="kt DENxSTF sites")

kt_RKxST_boxplot<-dep_pe_pg_99_filt_df %>% 
    filter(kinetochore==TRUE) %>%
    # select only tag columns, all replicates separately
    select(name, Sequence, all_of(grep("WT",pe_int_colnames,value=T))) %>% 
    # filter out proteins that have NA fc in all these columns
    # name in this case is actually a gene_name_site
    filter(if_any(-c(name,Sequence),~!is.na(.x))) %>%
    pivot_longer(-c(name,Sequence),names_to="sample",values_to="log2int") %>% 
    mutate(stage=ifelse(grepl("mitoM",sample),"mitoM",
                        ifelse(grepl("_MII_",sample),"MII",
                               ifelse(grepl("_MI_",sample),"MI","pro")))) %>%
    mutate(stage=factor(stage,levels=c("pro","MI","MII","mitoM"))) %>%
    mutate(replicate=ifelse(grepl("_1",sample),"rep1",
                            ifelse(grepl("_2",sample),"rep2","rep3"))) %>%
    mutate(int=2^log2int) %>%
    mutate(motif=grepl(".....[RK].[ST].......",Sequence)) %>%
    filter(motif==TRUE) %>%
    group_by(name, stage) %>%  # Group by gene.name.site and stage
    summarize(median_int = median(int, na.rm = TRUE), .groups = "drop") %>%
    ggplot(aes(x=stage,y=log10(median_int),fill=stage))+
    geom_boxplot(alpha=0.5)+
    stat_summary(fun.data=n_fun,geom="text")+
    stat_pwc(method="wilcox_test",label="p.adj.format",p.adjust.method = "bonferroni")+
    #geom_jitter(aes(color=replicate), size=1, alpha=0.9) +
    theme_classic()+
    theme(axis.text.x = element_text(angle=90,vjust=0.6),
          legend.position="none")+
    labs(title="kt RKxST sites")

kt_STSTDE_boxplot<-dep_pe_pg_99_filt_df %>% 
    filter(kinetochore==TRUE) %>%
    # select only tag columns, all replicates separately
    select(name, Sequence, all_of(grep("WT",pe_int_colnames,value=T))) %>% 
    # filter out proteins that have NA fc in all these columns
    # name in this case is actually a gene_name_site
    filter(if_any(-c(name,Sequence),~!is.na(.x))) %>%
    pivot_longer(-c(name,Sequence),names_to="sample",values_to="log2int") %>% 
    mutate(stage=ifelse(grepl("mitoM",sample),"mitoM",
                        ifelse(grepl("_MII_",sample),"MII",
                               ifelse(grepl("_MI_",sample),"MI","pro")))) %>%
    mutate(stage=factor(stage,levels=c("pro","MI","MII","mitoM"))) %>%
    mutate(replicate=ifelse(grepl("_1",sample),"rep1",
                            ifelse(grepl("_2",sample),"rep2","rep3"))) %>%
    mutate(int=2^log2int) %>%
    mutate(motif=grepl(".......[ST][STDE]......",Sequence)) %>%
    filter(motif==TRUE) %>%
    group_by(name, stage) %>%  # Group by gene.name.site and stage
    summarize(median_int = median(int, na.rm = TRUE), .groups = "drop") %>%
    ggplot(aes(x=stage,y=log10(median_int),fill=stage))+
    geom_boxplot(alpha=0.5)+
    stat_summary(fun.data=n_fun,geom="text")+
    stat_pwc(method="wilcox_test",label="p.adj.format",p.adjust.method = "bonferroni")+
    #geom_jitter(aes(color=replicate), size=1, alpha=0.9) +
    theme_classic()+
    theme(axis.text.x = element_text(angle=90,vjust=0.6),
          legend.position="none")+
    labs(title="kt STSTDE sites")

print_plot(ggarrange(kt_sites_boxplot,kt_SP_boxplot,
                     kt_DEN_boxplot,ncol=1),
           "kt_motif_sites_boxplot.pdf",3,12)

# NEW
print_plot(ggarrange(kt_SP_boxplot,kt_SPxKR_boxplot,
                     kt_DEN_boxplot,kt_DENF_boxplot,
                     kt_RKxST_boxplot,kt_STSTDE_boxplot,ncol=3),
           "new_kt_motif_sites_boxplot.pdf",12,6)

# combine all 6 boxplots

print_plot(ggarrange(pe_total_phos_boxplot+theme(legend.position="none")+
                         stat_pwc(method="wilcox_test",label="p.adj.format",p.adjust.method = "bonferroni"),all_SP_boxplot,
                     all_DEN_boxplot,kt_sites_boxplot,kt_SP_boxplot,
                     kt_DEN_boxplot,nrow=2,ncol=3),
           "motif_sites_boxplot.pdf",12,12)

# heatmap of individual core [ST]P and [DEN]x[ST] sites (IDs)
# heatmap of motif sites ####

core_kt_polo_motif_heatmap<-dep_pe_pg_99_filt_df %>% 
    filter(new_cat %in% c("cbf1_3","inner_kt","KMN","dam1")) %>%
    # select only tag columns, all replicates separately
    select(name, Sequence, all_of(grep("WT",pe_int_colnames,value=T))) %>% 
    # filter out proteins that have NA fc in all these columns
    # name in this case is actually a gene_name_site
    filter(if_any(-c(name,Sequence),~!is.na(.x))) %>%
    pivot_longer(-c(name,Sequence),names_to="sample",values_to="log2int") %>% 
    mutate(stage=ifelse(grepl("mitoM",sample),"mitoM",
                        ifelse(grepl("_MII_",sample),"MII",
                               ifelse(grepl("_MI_",sample),"MI","pro")))) %>%
    mutate(stage=factor(stage,levels=c("pro","MI","MII","mitoM"))) %>%
    mutate(replicate=ifelse(grepl("_1",sample),"rep1",
                            ifelse(grepl("_2",sample),"rep2","rep3"))) %>%
    mutate(int=2^log2int) %>%
    mutate(motif=grepl(".....[DEN].[ST].......",Sequence)) %>%
    filter(motif==TRUE) %>%
    # need to somehow summarize the 3 reps of each stage
    group_by(name,stage) %>%
    mutate(median_int=median(int,na.rm=T)) %>%
    # reorder names so they are ordered by int
    ungroup %>%
    group_by(name) %>%
    mutate(med_sum=sum(median_int,na.rm=T)) %>%
    ungroup() %>% 
    mutate(name=fct_reorder(name,med_sum)) %>%
    # mutate(detected=!is.na(median_int)) %>%
    # filter(detected==TRUE) %>%
    ggplot(aes(x=stage,y=name,fill=log10(median_int)))+
    geom_tile(color="black")+
    scale_fill_gradient(low="white",high="darkred")+
    theme_minimal()+
    theme(axis.text.x = element_text(angle=90,vjust=0.5))+
    labs(x="",y="",fill="log10int",title="Core KT [DEN]x[ST]* phospho-sites")

core_kt_SP_motif_heatmap<-dep_pe_pg_99_filt_df %>% 
    filter(new_cat %in% c("cbf1_3","inner_kt","KMN","dam1")) %>%
    # select only tag columns, all replicates separately
    select(name, Sequence, all_of(grep("WT",pe_int_colnames,value=T))) %>% 
    # filter out proteins that have NA fc in all these columns
    # name in this case is actually a gene_name_site
    filter(if_any(-c(name,Sequence),~!is.na(.x))) %>%
    pivot_longer(-c(name,Sequence),names_to="sample",values_to="log2int") %>% 
    mutate(stage=ifelse(grepl("mitoM",sample),"mitoM",
                        ifelse(grepl("_MII_",sample),"MII",
                               ifelse(grepl("_MI_",sample),"MI","pro")))) %>%
    mutate(stage=factor(stage,levels=c("pro","MI","MII","mitoM"))) %>%
    mutate(replicate=ifelse(grepl("_1",sample),"rep1",
                            ifelse(grepl("_2",sample),"rep2","rep3"))) %>%
    mutate(int=2^log2int) %>%
    mutate(motif=grepl(".......[ST]P......",Sequence)) %>%
    filter(motif==TRUE) %>%
    # need to somehow summarize the 3 reps of each stage
    group_by(name,stage) %>%
    mutate(median_int=median(int,na.rm=T)) %>%
    # reorder names so they are ordered by int
    ungroup %>%
    group_by(name) %>%
    mutate(med_sum=sum(median_int,na.rm=T)) %>%
    ungroup() %>% 
    mutate(name=fct_reorder(name,med_sum)) %>%
    ggplot(aes(x=stage,y=name,fill=log10(median_int)))+
    geom_tile(color="black")+
    scale_fill_gradient(low="white",high="darkorchid4")+
    theme_minimal()+
    theme(axis.text.x = element_text(angle=90,vjust=0.5))+
    labs(x="",y="",fill="log10int",title="Core KT [ST]*P phospho-sites")

print_plot(ggarrange(core_kt_polo_motif_heatmap,
                     core_kt_SP_motif_heatmap),
           "core_kt_motif_sites_heatmap.pdf",10,10)

# print_plot(ggarrange(core_kt_polo_motif_heatmap2,
#                      core_kt_SP_motif_heatmap),
#            "core_kt_motif_sites_heatmap2.pdf",10,10)

# protein subcomplex boxplots ####

# # add new cat column
dsn1_dep_df<-dsn1_dep_df %>%
    mutate(new_cat=ifelse(name %in% cbf1_3, "cbf1_3",
                          ifelse(name %in% inner_kt, "inner_kt",
                                 ifelse(name %in% KMN, "KMN",
                                        ifelse(name %in% dam1,"dam1",
                                               ifelse(name %in% accessory, "accessory_kt",
                                                      ifelse(name %in% maps, "maps_kt",
                                                             ifelse(name %in% cpc_sac,"cpc_sac",
                                                                   "other"))))))))
                                                             

all_prot_boxplot<-dsn1_dep_df %>% 
    select(name, new_cat, all_of(grep("WT",pg_colnames,value=T))) %>% 
    # filter out proteins that have NA fc in all these columns
    filter(if_any(-c(name,new_cat),~!is.na(.x))) %>%
    pivot_longer(-c(name,new_cat),names_to="sample",values_to="int") %>% 
    mutate(stage=ifelse(grepl("mitoM",sample),"mitoM",
                        ifelse(grepl("_MII_",sample),"MII",
                               ifelse(grepl("_MI_",sample),"MI","pro")))) %>%
    mutate(stage=factor(stage,levels=c("pro","MI","MII","mitoM"))) %>%
    mutate(replicate=ifelse(grepl("_1",sample),"rep1",
                            ifelse(grepl("_2",sample),"rep2","rep3"))) %>%
    # you must keep the below
    # otherwise the stat_summary count number
    # does not work
    mutate(int=2^int) %>%
    group_by(name, stage) %>%  # Group by name and stage
    summarize(median_int = median(int, na.rm = TRUE),.groups="drop") %>%
    ggplot(aes(x=stage,y=log10(median_int),fill=stage))+
    geom_boxplot(alpha=0.5)+
    stat_summary(fun.data=n_fun,geom="text")+
    stat_pwc(method="wilcox_test",label="p.adj.format",p.adjust.method = "bonferroni")+
    theme_classic()+
    theme(axis.text.x = element_text(angle=90,vjust=0.6),
          legend.position="none")+
    labs(title="all protein intensity")

print_plot(all_prot_boxplot,"all_prot_boxplot.pdf",6,8)


# create col counting whether protein is in the kinetochore list

dsn1_dep_df<-dsn1_dep_df %>% 
mutate(kinetochore=ifelse(Genes %in% master_kt_df$Gene,T,F))

# dep_pg_df<-dep_pg_df %>%
#     mutate(new_cat=ifelse(name %in% cbf1_3, "cbf1_3",
#                       ifelse(name %in% inner_kt, "inner_kt",
#                              ifelse(name %in% KMN, "KMN",
#                                     ifelse(name %in% dam1,"dam1",
#                                            ifelse(name %in% accessory, "accessory_kt",
#                                                   ifelse(name %in% maps, "maps_kt",
#                                                          ifelse(name %in% cpc_sac,"cpc_sac",
#                                                                 "other")))))))) 


pg_kt_subgroup_boxplot<-function(subgroup,plot_title){
    dsn1_dep_df %>% 
        # filter to one sub_complex at a time otherwise the plot is too large
        filter(new_cat==subgroup) %>%
        # select only tag columns, all replicates separately
        select(name, new_cat, all_of(grep("WT",pg_colnames,value=T))) %>% 
        # filter out proteins that have NA fc in all these columns
        # name in this case is actually a gene_name_site
        filter(if_any(-c(name,new_cat),~!is.na(.x))) %>%
        pivot_longer(-c(name,new_cat),names_to="sample",values_to="log2int") %>% 
        mutate(stage=ifelse(grepl("mitoM",sample),"mitoM",
                            ifelse(grepl("_MII_",sample),"MII",
                                   ifelse(grepl("_MI_",sample),"MI","pro")))) %>%
        mutate(stage=factor(stage,levels=c("pro","MI","MII","mitoM"))) %>%
        mutate(replicate=ifelse(grepl("_1",sample),"rep1",
                                ifelse(grepl("_2",sample),"rep2","rep3"))) %>%
        mutate(int=2^log2int) %>%
        group_by(name, stage) %>%  # Group by gene.name.site and stage
        summarize(median_int = median(int, na.rm = TRUE), .groups = "drop") %>%
        ggplot(aes(x=stage,y=log10(median_int),fill=stage))+
        geom_boxplot(alpha=0.5)+
        stat_summary(fun.data=n_fun,geom="text")+
        stat_pwc(method="wilcox_test",label="p.adj.format",p.adjust.method = "bonferroni")+
        #stat_summary(fun.data=n_fun,geom="boxplot",alpha=0.9)+
        #geom_jitter(aes(color=replicate), size=1, alpha=0.9) +
        geom_jitter(size=0.5, alpha=0.9) +
        theme_classic()+
        theme(axis.text.x = element_text(angle=90,vjust=0.6),
              legend.position="none")+
        labs(title=plot_title)
}

pg_kt_cbf3_boxplot<-pg_kt_subgroup_boxplot("cbf1_3","Cbf3")
pg_kt_innerkt_boxplot<-pg_kt_subgroup_boxplot("inner_kt","CCAN")
pg_kt_kmn_boxplot<-pg_kt_subgroup_boxplot("KMN","KMN")
pg_kt_dam1_boxplot<-pg_kt_subgroup_boxplot("dam1","Dam1c")
pg_kt_acc_boxplot<-pg_kt_subgroup_boxplot("accessory_kt","Accessory KT")
pg_kt_cpc_sac_boxplot<-pg_kt_subgroup_boxplot("cpc_sac","CPC/SAC")
pg_kt_maps_boxplot<-pg_kt_subgroup_boxplot("maps_kt","MAPs")

# revised function to make subgroup boxplot without stat_pwc
# (so axis is better scaled in absence of stats bars)
pg_kt_subgroup_boxplot_nostats<-function(subgroup,plot_title){
    dsn1_dep_df %>% 
        # filter to one sub_complex at a time otherwise the plot is too large
        filter(new_cat==subgroup) %>%
        # select only tag columns, all replicates separately
        select(name, new_cat, all_of(grep("WT",pg_colnames,value=T))) %>% 
        # filter out proteins that have NA fc in all these columns
        # name in this case is actually a gene_name_site
        filter(if_any(-c(name,new_cat),~!is.na(.x))) %>%
        pivot_longer(-c(name,new_cat),names_to="sample",values_to="log2int") %>% 
        mutate(stage=ifelse(grepl("mitoM",sample),"mitoM",
                            ifelse(grepl("_MII_",sample),"MII",
                                   ifelse(grepl("_MI_",sample),"MI","pro")))) %>%
        mutate(stage=factor(stage,levels=c("pro","MI","MII","mitoM"))) %>%
        mutate(replicate=ifelse(grepl("_1",sample),"rep1",
                                ifelse(grepl("_2",sample),"rep2","rep3"))) %>%
        mutate(int=2^log2int) %>%
        group_by(name, stage) %>%  # Group by gene.name.site and stage
        summarize(median_int = median(int, na.rm = TRUE), .groups = "drop") %>%
        ggplot(aes(x=stage,y=log10(median_int),fill=stage))+
        geom_boxplot(alpha=0.5)+
        stat_summary(fun.data=n_fun,geom="text")+
        #stat_pwc(method="wilcox_test",label="p.adj.format",p.adjust.method = "bonferroni")+
        #stat_summary(fun.data=n_fun,geom="boxplot",alpha=0.9)+
        #geom_jitter(aes(color=replicate), size=1, alpha=0.9) +
        geom_jitter(size=0.5, alpha=0.9) +
        theme_classic()+
        theme(axis.text.x = element_text(angle=90,vjust=0.6),
              legend.position="none")+
        labs(title=plot_title)
}

pg_kt_cbf3_boxplot_nostat<-pg_kt_subgroup_boxplot_nostats("cbf1_3","Cbf3")
pg_kt_innerkt_boxplot_nostat<-pg_kt_subgroup_boxplot_nostats("inner_kt","CCAN")
# pg_kt_kmn_boxplot<-pg_kt_subgroup_boxplot("KMN","KMN")
# pg_kt_dam1_boxplot<-pg_kt_subgroup_boxplot("dam1","Dam1c")
# pg_kt_acc_boxplot<-pg_kt_subgroup_boxplot("accessory_kt","Accessory KT")
pg_kt_cpc_sac_boxplot_nostat<-pg_kt_subgroup_boxplot_nostats("cpc_sac","CPC/SAC")
#pg_kt_maps_boxplot<-pg_kt_subgroup_boxplot("maps_kt","MAPs")


# all KT proteins boxplot (to make even 8 plots)
all_pg_kt_boxplot<-dsn1_dep_df %>% 
    # filter to one sub_complex at a time otherwise the plot is too large
    filter(kinetochore==TRUE) %>%
    # select only tag columns, all replicates separately
    select(name, kinetochore, all_of(grep("WT",pg_colnames,value=T))) %>% 
    # filter out proteins that have NA fc in all these columns
    # name in this case is actually a gene_name_site
    filter(if_any(-c(name,kinetochore),~!is.na(.x))) %>%
    pivot_longer(-c(name,kinetochore),names_to="sample",values_to="log2int") %>% 
    mutate(stage=ifelse(grepl("mitoM",sample),"mitoM",
                        ifelse(grepl("_MII_",sample),"MII",
                               ifelse(grepl("_MI_",sample),"MI","pro")))) %>%
    mutate(stage=factor(stage,levels=c("pro","MI","MII","mitoM"))) %>%
    mutate(replicate=ifelse(grepl("_1",sample),"rep1",
                            ifelse(grepl("_2",sample),"rep2","rep3"))) %>%
    mutate(int=2^log2int) %>%
    group_by(name, stage) %>%  # Group by gene.name.site and stage
    summarize(median_int = median(int, na.rm = TRUE), .groups = "drop") %>%
    ggplot(aes(x=stage,y=log10(median_int),fill=stage))+
    geom_boxplot(alpha=0.5)+
    stat_summary(fun.data=n_fun,geom="text")+
    stat_pwc(method="wilcox_test",label="p.adj.format",p.adjust.method = "bonferroni")+
    #stat_summary(fun.data=n_fun,geom="boxplot",alpha=0.9)+
    #geom_jitter(aes(color=replicate), size=1, alpha=0.9) +
    geom_jitter(size=0.5, alpha=0.9) +
    theme_classic()+
    theme(axis.text.x = element_text(angle=90,vjust=0.6),
          legend.position="none")+
    labs(title="all kt proteins")

# no stats all kT proteins
# all KT proteins boxplot (to make even 8 plots)
all_pg_kt_boxplot_nostats<-dsn1_dep_df %>% 
    # filter to one sub_complex at a time otherwise the plot is too large
    filter(kinetochore==TRUE) %>%
    # select only tag columns, all replicates separately
    select(name, kinetochore, all_of(grep("WT",pg_colnames,value=T))) %>% 
    # filter out proteins that have NA fc in all these columns
    # name in this case is actually a gene_name_site
    filter(if_any(-c(name,kinetochore),~!is.na(.x))) %>%
    pivot_longer(-c(name,kinetochore),names_to="sample",values_to="log2int") %>% 
    mutate(stage=ifelse(grepl("mitoM",sample),"mitoM",
                        ifelse(grepl("_MII_",sample),"MII",
                               ifelse(grepl("_MI_",sample),"MI","pro")))) %>%
    mutate(stage=factor(stage,levels=c("pro","MI","MII","mitoM"))) %>%
    mutate(replicate=ifelse(grepl("_1",sample),"rep1",
                            ifelse(grepl("_2",sample),"rep2","rep3"))) %>%
    mutate(int=2^log2int) %>%
    group_by(name, stage) %>%  # Group by gene.name.site and stage
    summarize(median_int = median(int, na.rm = TRUE), .groups = "drop") %>%
    ggplot(aes(x=stage,y=log10(median_int),fill=stage))+
    geom_boxplot(alpha=0.5)+
    stat_summary(fun.data=n_fun,geom="text")+
    #stat_pwc(method="wilcox_test",label="p.adj.format",p.adjust.method = "bonferroni")+
    #stat_summary(fun.data=n_fun,geom="boxplot",alpha=0.9)+
    #geom_jitter(aes(color=replicate), size=1, alpha=0.9) +
    geom_jitter(size=0.5, alpha=0.9) +
    theme_classic()+
    theme(axis.text.x = element_text(angle=90,vjust=0.6),
          legend.position="none")+
    labs(title="all kt proteins")


pg_kt_boxplots<-ggarrange(all_pg_kt_boxplot,pg_kt_cbf3_boxplot,pg_kt_innerkt_boxplot,pg_kt_kmn_boxplot,
                       pg_kt_dam1_boxplot,pg_kt_acc_boxplot,pg_kt_cpc_sac_boxplot,
                       pg_kt_maps_boxplot,nrow=2,ncol=4,align="h")

print_plot(pg_kt_boxplots,"pg_kt_boxplots.pdf",12,8)

# revised so plots with no sig diffs do not have
# wider axis to allow for stats bars
# only plots with sigdiffs are : KMN, Dam1, Accessory,MAPs

pg_kt_boxplots_revised<-ggarrange(all_pg_kt_boxplot_nostats,pg_kt_cbf3_boxplot_nostat,pg_kt_innerkt_boxplot_nostat,pg_kt_kmn_boxplot,
                          pg_kt_dam1_boxplot,pg_kt_acc_boxplot,pg_kt_cpc_sac_boxplot_nostat,
                          pg_kt_maps_boxplot,nrow=2,ncol=4,align="h")

print_plot(pg_kt_boxplots_revised,"pg_kt_boxplots_revised.pdf",12,8)

# protein dynamic boxplot ####

prot_dynamic_boxplot<-dsn1_dep_df %>% 
    filter(kinetochore==T) %>%
    #filter(Gene.Names=="CTF13") %>%
    # only select tag columns which have "WT" in their name
    select(name, all_of(grep("WT",pg_colnames,value=T))) %>% 
    # filter out proteins that have NA in all these columns
    filter(if_any(-c(name),~!is.na(.x))) %>%
    pivot_longer(-c(name),names_to="sample",values_to="log2int") %>% 
    mutate(stage=ifelse(grepl("mitoM",sample),"mitoM",
                        ifelse(grepl("_MII_",sample),"MII",
                               ifelse(grepl("_MI_",sample),"MI","pro")))) %>%
    mutate(stage=factor(stage,levels=c("pro","MI","MII","mitoM"))) %>%
    mutate(replicate=ifelse(grepl("_1",sample),"rep1",
                            ifelse(grepl("_2",sample),"rep2","rep3"))) %>%
    group_by(name,stage) %>%
    summarize(stage_median=median(log2int,na.rm=T)) %>%
    ungroup() %>%
    group_by(name) %>%
    mutate(stage_range=max(stage_median,na.rm=T)-min(stage_median,na.rm=T)) %>%
    ungroup() %>% 
    mutate(name=paste0(name," (",round(stage_range,1),")")) %>%
    mutate(name=as.factor(name)) %>%
    # mutate(stage_range=max(stage_median,na.rm=T)-min(stage_median,na.rm=T)) %>%
    # mutate(name=fct_reorder(name,stage_range,na.rm=T)) %>%
     mutate(name=fct_reorder(name,stage_range,na.rm=T)) %>%
    ggplot(aes(x=name,y=stage_median))+
    geom_boxplot()+
    geom_jitter(aes(color=stage),alpha=1)+
    #coord_flip()+
    labs(x="",y="log2(dsn1-scaled abundance)")+
    theme_classic()+
    theme(axis.text.y=element_text(size=rel(1.5)),axis.text.x=element_text(angle=90,vjust=0.5))

print_plot(prot_dynamic_boxplot,"prot_dynamic_boxplot.pdf",15,6)

# KT proteins int boxplot

kt_prot_boxplot<-dsn1_dep_df %>% 
    # filter to KT proteins only
    filter(name %in% master_kt_df$Gene) %>%
    select(name, new_cat, all_of(grep("WT",pg_colnames,value=T))) %>% 
    # filter out proteins that have NA fc in all these columns
    filter(if_any(-c(name,new_cat),~!is.na(.x))) %>%
    pivot_longer(-c(name,new_cat),names_to="sample",values_to="int") %>% 
    mutate(stage=ifelse(grepl("mitoM",sample),"mitoM",
                        ifelse(grepl("_MII_",sample),"MII",
                               ifelse(grepl("_MI_",sample),"MI","pro")))) %>%
    mutate(stage=factor(stage,levels=c("pro","MI","MII","mitoM"))) %>%
    mutate(replicate=ifelse(grepl("_1",sample),"rep1",
                            ifelse(grepl("_2",sample),"rep2","rep3"))) %>%
    # you must keep the below
    # otherwise the stat_summary count number
    # does not work
    mutate(int=2^int) %>%
    group_by(name, stage) %>%  # Group by name and stage
    summarize(median_int = median(int, na.rm = TRUE),.groups="drop") %>%
    ggplot(aes(x=stage,y=log10(median_int),fill=stage))+
    geom_boxplot(alpha=0.5)+
    stat_summary(fun.data=n_fun,geom="text")+
    stat_pwc(method="wilcox_test",label="p.adj.format",p.adjust.method = "bonferroni")+
    theme_classic()+
    theme(axis.text.x = element_text(angle=90,vjust=0.6),
          legend.position="none")+
    labs(title="kt protein intensity")

print_plot(kt_prot_boxplot,"kt_prot_boxplot.pdf",6,8)

# new GO section ####

# each stage vs the other 3 stages
# of sigdiff proteins (called by DEP)
# what is median fc for that stage vs other 3 (make sure directions)

diff_cols<-dsn1_dep_df %>% 
    select(grep("_diff",colnames(.),value=T)) %>% colnames()

# start with pro
#grep("pro",diff_cols,value=T)
# one needs to be flipped in direction
# WT_MI_vs_WT_pro_diff
# WT_MII_vs_WT_pro_diff
# WT_pro_vs_WT_mitoM_diff

# of the proteins most diff 
# and which are not KT proteins
# what are these proteins
# create combined list for the 3 comparisons

# always make df with _diffs arranged in same stage order 
# for consistency:
# pro -> MI -> MII -> mitoM
# if you start in the middle of the seq go back to the beginning?

pro_enriched_genes<-bind_rows(dsn1_dep_df %>%
    select(name,WT_MI_vs_WT_pro_diff) %>%
    # the negative values here are proteins higher in pro
    arrange((WT_MI_vs_WT_pro_diff)) %>%
    #head() # ZIP1, NDJ1 are most diff higher in prophase vs MI
    # filter out kinetochore proteins
    #nrow()
    filter(!name %in% master_kt_df$Gene) %>% #nrow()
    # take the top 50 proteins
    head(50) %>% select(name),
dsn1_dep_df %>%
    select(name,WT_MII_vs_WT_pro_diff) %>%
    # the negative values here are proteins higher in pro
    arrange((WT_MII_vs_WT_pro_diff)) %>%
    #head() # ZIP1, NDJ1 are most diff higher in prophase vs MII
    # filter out kinetochore proteins
    #nrow()
    filter(!name %in% master_kt_df$Gene) %>% #nrow()
    # take the top 50 proteins
    head(50) %>% select(name),
dsn1_dep_df %>%
    select(name,WT_pro_vs_WT_mitoM_diff) %>%
    # the POSITIVE values here are proteins higher in pro
    arrange(desc(WT_pro_vs_WT_mitoM_diff)) %>%
    #head() # ADY2,MDH2 are most diff higher in prophase vs mitoM
    # filter out kinetochore proteins
    #nrow()
    filter(!name %in% master_kt_df$Gene) %>% #nrow()
    # take the top 50 proteins
    head(50) %>% select(name)) %>%
    distinct() %>%
    unlist() %>% unname()

library(gprofiler2)

pro_enriched_result<-gost(pro_enriched_genes,organism="scerevisiae",significant = T,
     evcodes=T)$result 

# copy 3 top20 gene lists as df to excel
dsn1_dep_df %>%
              select(name,WT_MI_vs_WT_pro_diff) %>%
              # the negative values here are proteins higher in pro
              arrange((WT_MI_vs_WT_pro_diff)) %>%
              #head() # ZIP1, NDJ1 are most diff higher in prophase vs MI
              # filter out kinetochore proteins
              #nrow()
              filter(!name %in% master_kt_df$Gene) %>% #nrow()
              # take the top 20 proteins
              head(20) %>% select(name) %>% write_clip()

dsn1_dep_df %>%
              select(name,WT_MII_vs_WT_pro_diff) %>%
              # the negative values here are proteins higher in pro
              arrange((WT_MII_vs_WT_pro_diff)) %>%
              #head() # ZIP1, NDJ1 are most diff higher in prophase vs MII
              # filter out kinetochore proteins
              #nrow()
              filter(!name %in% master_kt_df$Gene) %>% #nrow()
              # take the top 20 proteins
              head(20) %>% select(name) %>% write_clip()

dsn1_dep_df %>%
              select(name,WT_pro_vs_WT_mitoM_diff) %>%
              # the POSITIVE values here are proteins higher in pro
              arrange(desc(WT_pro_vs_WT_mitoM_diff)) %>%
              #head() # ADY2,MDH2 are most diff higher in prophase vs mitoM
              # filter out kinetochore proteins
              #nrow()
              filter(!name %in% master_kt_df$Gene) %>% #nrow()
              # take the top 20 proteins
              head(20) %>% select(name) %>% write_clip()

# copy_GO_table<-function(dataframe){
#     dataframe %>%
#         select(term_name,term_id,p_value,source) %>%
#         write_clip()
# }

# view top GO terms
pro_enriched_result %>%
    select(term_name,term_id,p_value) %>%
    head(20)
#GO:0140527 reciprocal homologous recombination
#GO:0007129 homologous chromosome pairing at meiosis
#GO:0045333 cellular respiration

# MI genes

grep("_MI_",diff_cols,value=T)

MI_enriched_genes<-bind_rows(dsn1_dep_df %>%
    select(name,WT_MII_vs_WT_MI_diff) %>%
    # the NEGATIVE values here are proteins higher in MI vs MII
    arrange((WT_MII_vs_WT_MI_diff)) %>%
    #head() # SGO1,DBF4 are most diff higher in MI vs MII
    # filter out kinetochore proteins
    #nrow()
    filter(!name %in% master_kt_df$Gene) %>% #nrow()
    # take the top 50 proteins
    head(50) %>% select(name),
dsn1_dep_df %>%
    select(name,WT_MI_vs_WT_mitoM_diff) %>%
    # the positive values here are proteins higher in MI
    arrange(desc(WT_MI_vs_WT_mitoM_diff)) %>%
    #head() # ADY2,CIT3 are most diff higher in MI vs mitoM
    # filter out kinetochore proteins
    #nrow()
    filter(!name %in% master_kt_df$Gene) %>% #nrow()
    # take the top 50 proteins
    head(50) %>% select(name),
dsn1_dep_df %>%
    select(name,WT_MI_vs_WT_pro_diff) %>%
    # the positive values here are proteins higher in MI
    arrange(desc(WT_MI_vs_WT_pro_diff)) %>%
    #head() # DAD1,DAD3 are most diff higher in MI vs pro
    # filter out kinetochore proteins
    #nrow()
    filter(!name %in% master_kt_df$Gene) %>% #nrow()
    # take the top 50 proteins
    head(50) %>% select(name)) %>%
    distinct() %>%
    unlist() %>% unname()

MI_enriched_result<-gost(MI_enriched_genes,organism="scerevisiae",significant = T,
                          evcodes=T)$result 

MI_enriched_result %>%
    select(term_name,term_id,p_value) %>%
    head(20)
#GO:0007051 spindle organization
#GO:0051300 spindle pole body organization
#GO:0051276 chromosome organization

# copy 3 top20 gene lists to excel
dsn1_dep_df %>%
              select(name,WT_MII_vs_WT_MI_diff) %>%
              # the NEGATIVE values here are proteins higher in MI vs MII
              arrange((WT_MII_vs_WT_MI_diff)) %>%
              #head() # SGO1,DBF4 are most diff higher in MI vs MII
              # filter out kinetochore proteins
              #nrow()
              filter(!name %in% master_kt_df$Gene) %>% #nrow()
              # take the top 50 proteins
              head(20) %>% select(name) %>% write_clip()

dsn1_dep_df %>%
              select(name,WT_MI_vs_WT_mitoM_diff) %>%
              # the positive values here are proteins higher in MI
              arrange(desc(WT_MI_vs_WT_mitoM_diff)) %>%
              #head() # ADY2,CIT3 are most diff higher in MI vs mitoM
              # filter out kinetochore proteins
              #nrow()
              filter(!name %in% master_kt_df$Gene) %>% #nrow()
              # take the top 20 proteins
              head(20) %>% select(name) %>% write_clip()

dsn1_dep_df %>%
              select(name,WT_MI_vs_WT_pro_diff) %>%
              # the positive values here are proteins higher in MI
              arrange(desc(WT_MI_vs_WT_pro_diff)) %>%
              #head() # DAD1,DAD3 are most diff higher in MI vs pro
              # filter out kinetochore proteins
              #nrow()
              filter(!name %in% master_kt_df$Gene) %>% #nrow()
              # take the top 20 proteins
              head(20) %>% select(name) %>% write_clip()

# MII genes

grep("_MII_",diff_cols,value=T)

MII_enriched_genes<-bind_rows(dsn1_dep_df %>%
    select(name,WT_MII_vs_WT_MI_diff) %>%
    # the positive values here are proteins higher in MII
    arrange(desc(WT_MII_vs_WT_MI_diff)) %>%
    #head() # SPS1,CDC15 are most diff higher in MII vs MI
    # filter out kinetochore proteins
    #nrow()
    filter(!name %in% master_kt_df$Gene) %>% #nrow()
    # take the top 50 proteins
    head(50) %>% select(name),
dsn1_dep_df %>%
    select(name,WT_MII_vs_WT_mitoM_diff) %>%
    # the positive values here are proteins higher in MII
    arrange(desc(WT_MII_vs_WT_mitoM_diff)) %>%
    #head() # ADY2,YAT2 are most diff higher in MII vs mitoM
    # filter out kinetochore proteins
    #nrow()
    filter(!name %in% master_kt_df$Gene) %>% #nrow()
    # take the top 50 proteins
    head(50) %>% select(name),
dsn1_dep_df %>%
    select(name,WT_MII_vs_WT_pro_diff) %>%
    # the positive values here are proteins higher in MII
    arrange(desc(WT_MII_vs_WT_pro_diff)) %>%
    #head() # DAD1,FIN1 are most diff higher in MII vs pro
    # filter out kinetochore proteins
    #nrow()
    filter(!name %in% master_kt_df$Gene) %>% #nrow()
    # take the top 50 proteins
    head(50) %>% select(name)) %>%
    distinct() %>% unlist() %>% unname()

MII_enriched_result<-gost(MII_enriched_genes,organism="scerevisiae",significant = T,
                         evcodes=T)$result 

MII_enriched_result %>%
    select(term_name,term_id,p_value) %>%
    head(30)
#GO:0043934 sporulation
#GO:0022402 cell cycle process
#GO:0042244 spore wall assembly

# copy 3 top20 gene lists to excel
dsn1_dep_df %>%
    select(name,WT_MII_vs_WT_MI_diff) %>%
    # the positive values here are proteins higher in MII
    arrange(desc(WT_MII_vs_WT_MI_diff)) %>%
    #head() # SPS1,CDC15 are most diff higher in MII vs MI
    # filter out kinetochore proteins
    #nrow()
    filter(!name %in% master_kt_df$Gene) %>% #nrow()
    # take the top 20 proteins
    head(20) %>% select(name) %>% write_clip()

dsn1_dep_df %>%
    select(name,WT_MII_vs_WT_mitoM_diff) %>%
    # the positive values here are proteins higher in MII
    arrange(desc(WT_MII_vs_WT_mitoM_diff)) %>%
    #head() # ADY2,YAT2 are most diff higher in MII vs mitoM
    # filter out kinetochore proteins
    #nrow()
    filter(!name %in% master_kt_df$Gene) %>% #nrow()
    # take the top 20 proteins
    head(20) %>% select(name) %>% write_clip()

dsn1_dep_df %>%
    select(name,WT_MII_vs_WT_pro_diff) %>%
    # the positive values here are proteins higher in MII
    arrange(desc(WT_MII_vs_WT_pro_diff)) %>%
    #head() # DAD1,FIN1 are most diff higher in MII vs pro
    # filter out kinetochore proteins
    #nrow()
    filter(!name %in% master_kt_df$Gene) %>% #nrow()
    # take the top 20 proteins
    head(20) %>% select(name) %>% write_clip()

# mitoM genes

grep("_mitoM_",diff_cols,value=T)

mitoM_enriched_genes<-bind_rows(dsn1_dep_df %>%
    select(name,WT_MI_vs_WT_mitoM_diff) %>%
    # the negative values here are proteins higher in mitoM
    arrange((WT_MI_vs_WT_mitoM_diff)) %>%
    #head() # BUD20,UBA4 are most diff higher in mitoM vs MI
    # filter out kinetochore proteins
    #nrow()
    filter(!name %in% master_kt_df$Gene) %>% #nrow()
    # take the top 50 proteins
    head(50) %>% select(name),
dsn1_dep_df %>%
    select(name,WT_MII_vs_WT_mitoM_diff) %>%
    # the negative values here are proteins higher in mitoM
    arrange((WT_MII_vs_WT_mitoM_diff)) %>%
    #head() # UBA4,HXT2 are most diff higher in mitoM vs MII
    # filter out kinetochore proteins
    #nrow()
    filter(!name %in% master_kt_df$Gene) %>% #nrow()
    # take the top 50 proteins
    head(50) %>% select(name),
dsn1_dep_df %>%
    select(name,WT_pro_vs_WT_mitoM_diff) %>%
    # the negative values here are proteins higher in mitoM
    arrange((WT_pro_vs_WT_mitoM_diff)) %>%
    #head() # DAD1,BUD20 are most diff higher in mitoM vs pro
    # filter out kinetochore proteins
    #nrow()
    filter(!name %in% master_kt_df$Gene) %>% #nrow()
    # take the top 50 proteins
    head(50) %>% select(name)) %>%
    distinct() %>% unlist() %>% unname()

mitoM_enriched_result<-gost(mitoM_enriched_genes,organism="scerevisiae",significant = T,
                          evcodes=T)$result 

mitoM_enriched_result %>%
    select(term_name,term_id,p_value) %>% 
    head(20)
#GO:0042254 ribosome biogenesis
#GO:0005730 nucleolus
#GO:0031981 nuclear lumen
#only 29 terms aand they are all ribosome or membrane related.

# copy 3 top20 gene lists to excel
dsn1_dep_df %>%
    select(name,WT_MI_vs_WT_mitoM_diff) %>%
    # the negative values here are proteins higher in mitoM
    arrange((WT_MI_vs_WT_mitoM_diff)) %>%
    #head() # BUD20,UBA4 are most diff higher in mitoM vs MI
    # filter out kinetochore proteins
    #nrow()
    filter(!name %in% master_kt_df$Gene) %>% #nrow()
    # take the top 20 proteins
    head(20) %>% select(name) %>% write_clip()

dsn1_dep_df %>%
    select(name,WT_MII_vs_WT_mitoM_diff) %>%
    # the negative values here are proteins higher in mitoM
    arrange((WT_MII_vs_WT_mitoM_diff)) %>%
    #head() # UBA4,HXT2 are most diff higher in mitoM vs MII
    # filter out kinetochore proteins
    #nrow()
    filter(!name %in% master_kt_df$Gene) %>% #nrow()
    # take the top 20 proteins
    head(20) %>% select(name) %>% write_clip()

dsn1_dep_df %>%
    select(name,WT_pro_vs_WT_mitoM_diff) %>%
    # the negative values here are proteins higher in mitoM
    arrange((WT_pro_vs_WT_mitoM_diff)) %>%
    #head() # DAD1,BUD20 are most diff higher in mitoM vs pro
    # filter out kinetochore proteins
    #nrow()
    filter(!name %in% master_kt_df$Gene) %>% #nrow()
    # take the top 20 proteins
    head(20) %>% select(name) %>% write_clip()

# create combined p-value plot of 12 enriched terms.
# bar plot
# y axis is the terms
# x axis is the p-value (-log)
# can draw a line for 0.05
# what is the structure of the df needed
# need the following info
# terms, p-value, stage of enrichment (pro, MI etc)
    
enriched_df<-rbind(
pro_enriched_result %>%
    select(term_name,term_id,p_value) %>% 
    filter(term_id %in% c("GO:0140527","GO:0007129","GO:0045333")) %>%
    cbind(stage="pro"),

MI_enriched_result %>%
    select(term_name,term_id,p_value) %>% 
    filter(term_id %in% c("GO:0007051","GO:0007059","GO:0051276")) %>%
    cbind(stage="MI"),

MII_enriched_result %>%
    select(term_name,term_id,p_value) %>% 
    filter(term_id %in% c("GO:0043934","GO:0022402","GO:0042244")) %>%
    cbind(stage="MII"),

mitoM_enriched_result %>%
    select(term_name,term_id,p_value) %>% 
    filter(term_id %in% c("GO:0042254","GO:0005730","GO:0031981")) %>%
    cbind(stage="mitoM")) 

combi_GO_bar<-enriched_df %>%
    mutate(term=paste0(term_name," ",term_id)) %>%
    mutate(stage=factor(stage,levels=c("pro","MI","MII","mitoM"))) %>%
    mutate(log_pval = -log10(p_value)) %>%
    group_by(stage) %>%
    # Re-factor term_name within each stage so only 3 terms show up per facet
    mutate(term = factor(term, levels = term[order(log_pval)])) %>%
    ggplot(aes(x = log_pval, y = term, fill = stage)) +
    geom_bar(stat = "identity") +
    facet_wrap(~stage, scales = "free_y",ncol=1) +
    labs(
        x = expression(-log[10](p~value)),
        y = "",
        title = ""
    ) +
    theme_minimal() +
    theme(strip.text = element_text(face = "bold"),
          legend.position="none")+
    geom_vline(xintercept=-log10(0.05),linetype="dashed")
    
print_plot(combi_GO_bar,"combi_GO_bar.pdf",6,4)

# PROTEIN NUMBER BAR ####
# use dsn1_dep_df

prot_num_bar<-dsn1_dep_df %>%
    summarize(across(all_of(pg_colnames), ~sum(!is.na(.)))) %>%
    pivot_longer(cols=all_of(pg_colnames),names_to="sample",
                 values_to="number") %>%
    filter(grepl("WT",sample)) %>%
    mutate(sample=factor(sample,levels=c("WT_pro_1",
                                         "WT_pro_2",
                                         "WT_pro_3",
                                         "WT_MI_1",
                                         "WT_MI_2",
                                         "WT_MI_3",
                                         "WT_MII_1",
                                         "WT_MII_2",
                                         "WT_MII_3",
                                         "WT_mitoM_1",
                                         "WT_mitoM_2",
                                         "WT_mitoM_3"))) %>%
    ggplot(aes(x=number,y=sample,fill=sample))+
    geom_bar(stat="identity")+
    scale_fill_viridis_d()+
    theme_classic()+
    theme(legend.position="none")+
    labs(x="number",y="",title="Proteins")+
    scale_x_continuous(expand=expansion(mult=c(0,0.2)))+
    geom_text(aes(label=number), hjust=-0.2, color="black", size=4)  # Adding text labels

phos_num_bar<-dep_pe_pg_99_filt_df %>% 
    summarize(across(all_of(pe_int_colnames), ~sum(!is.na(.)))) %>%
    pivot_longer(cols=all_of(pe_int_colnames),names_to="sample",
                 values_to="number") %>%
    filter(grepl("WT",sample)) %>%
    mutate(sample=factor(sample,levels=c("WT_pro_1",
                                         "WT_pro_2",
                                         "WT_pro_3",
                                         "WT_MI_1",
                                         "WT_MI_2",
                                         "WT_MI_3",
                                         "WT_MII_1",
                                         "WT_MII_2",
                                         "WT_MII_3",
                                         "WT_mitoM_1",
                                         "WT_mitoM_2",
                                         "WT_mitoM_3"))) %>%
    ggplot(aes(x=number,y=sample,fill=sample))+
    geom_bar(stat="identity")+
    scale_fill_viridis_d()+
    theme_classic()+
    theme(legend.position="none")+
    labs(x="number",y="",title="Phospho-sites")+
    scale_x_continuous(expand=expansion(mult=c(0,0.2)))+
    geom_text(aes(label=number), hjust=-0.2, color="black", size=4)  # Adding text labels

print_plot(prot_num_bar,"prot_num_bar.pdf",5,3)
print_plot(phos_num_bar,"phos_num_bar.pdf",5,3)

# KT Protein HEATMAPS ####
# sub_groups heatmaps with all 4 stages

sub_heatmap<-function(subgroup,plot_title){
    dsn1_dep_df %>%
        filter(new_cat==subgroup) %>%
        select(name,pg_colnames) %>%
        pivot_longer(cols=-name,names_to="sample",values_to="int") %>%
        filter(grepl("WT",sample)) %>%
        mutate(stage=ifelse(grepl("pro",sample),"pro",
                            ifelse(grepl("_MI_",sample),"MI",
                                   ifelse(grepl("_MII_",sample),"MII",
                                          "mitoM")))) %>%
        mutate(stage = factor(stage, levels = c("pro", "MI", "MII", "mitoM"), ordered = TRUE)) %>%  # ✨ make ordered
        group_by(name,stage) %>%
        summarize(med_int=median(int,na.rm=T),.groups="drop") %>%
        # summarize(min_med_int = min(med_int, na.rm = TRUE),
        #           max_med_int = max(med_int, na.rm = TRUE))
        mutate(name=fct_reorder2(name,dplyr::desc(stage),dplyr::desc(med_int),.na_rm=F)) %>%
        # mutate(name=fct_reorder2(name,med_int,stage,.na_rm=F)) %>%
        mutate(med_int=2^med_int) %>%
        ggplot(aes(x=stage,y=name,fill=med_int))+
        geom_tile(color="black")+
        scale_fill_gradient(low="white",high="darkblue")+
        #scale_fill_distiller(palette="RdBu")+
        theme_minimal()+
        theme(axis.text.x = element_text(angle=90,vjust=0.5))+
        #coord_flip()+
        labs(title=plot_title,x="",y="")
}

print_plot(sub_heatmap("cbf1_3","Cbf3"),"cbf3_heat.pdf",3,1.5)
print_plot(sub_heatmap("maps_kt","MAPs"),"maps_heat.pdf",3,2)
print_plot(sub_heatmap("cpc_sac","CPC/SAC"),"cpc_heat.pdf",3.2,2.125)
print_plot(sub_heatmap("accessory_kt","Accessory"),"acc_heat.pdf",3.2,3)
print_plot(sub_heatmap("dam1","Dam1"),"dam1_heat.pdf",3.5,2.3)
print_plot(sub_heatmap("KMN","KMN"),"kmn_heat.pdf",3.5,2.3)
print_plot(sub_heatmap("inner_kt","CCAN"),"ccan_heat.pdf",3.5,3.2)


# how many proteins are phos by diff kinases in inner vs outer KT?
# inner v outer barplots ####

# create detected_df2

detected_df2<-dep_pe_pg_99_filt_df %>%
    filter(kinetochore==TRUE) %>%
    # select only tag columns, all replicates separately
    select(Gene.Names,new_cat,name, Sequence,all_of(grep("WT",pe_int_colnames,value=T))) %>%
    # filter out proteins that have NA in all these columns
    filter(if_any(-name,~!is.na(.x))) %>%
    pivot_longer(-c(Gene.Names,new_cat,name,Sequence),names_to="sample",values_to="log2int") %>%
    mutate(stage=ifelse(grepl("mitoM",sample),"mitoM",
                        ifelse(grepl("_MII_",sample),"MII",
                               ifelse(grepl("_MI_",sample),"MI","pro")))) %>%
    mutate(stage=factor(stage,levels=c("pro","MI","MII","mitoM"))) %>%
    mutate(replicate=ifelse(grepl("_1",sample),"rep1",
                            ifelse(grepl("_2",sample),"rep2","rep3"))) %>%
    mutate(int=2^log2int) %>%
    group_by(Gene.Names,new_cat,name, stage, Sequence) %>%  # Group by gene.name.site and stage
    summarize(median_int = median(int, na.rm = TRUE), .groups = "drop") %>%
    group_by(stage) %>%
    mutate(detected=!is.na(median_int)) %>%
    # head()
    # group_by(stage) %>%
    #     summarize(n=sum(!is.na(median_int))) #100,143,119,124 same numbers as boxplots
    select(Gene.Names, new_cat, name, stage,Sequence,detected)

STP_subcomplex_bar<-detected_df2 %>%
        filter(detected==TRUE) %>% 
        # create col that says whether seq matches motif or not
        mutate(motif=grepl(".......[ST]P......",Sequence)) %>%
        group_by(stage,new_cat) %>%
        summarize(
            total_sequences = n(),
            motif_sequences = sum(motif)
        )  %>%
        mutate(perc_match=motif_sequences/total_sequences*100) %>%
        mutate(stage=factor(stage,levels=c("pro","MI","MII","mitoM"))) %>%
        #select(stage,perc_match) %>% head(10)
        filter(new_cat %in% c("cbf1_3","inner_kt","KMN","dam1")) %>%
     # factor levels so that groups go from inner to outer kinetochore
        mutate(new_cat=factor(new_cat,levels= c("cbf1_3","inner_kt","KMN","dam1"))) %>%
# check numbers match numbers of sites at each stage shown in boxplots
# they all match.
    ggplot(aes(x=interaction(stage,new_cat),y=perc_match,fill=new_cat))+
        geom_bar(stat="identity",position="stack")+
        #coord_flip()+
        theme_classic()+
        labs(title="[ST]*P sites",y="percent_match",x="")+
        scale_y_continuous(expand=expansion(mult=c(0,0.2)))+
        theme(legend.position="none",axis.text.x = element_text(angle=90,vjust=0.5))+
    geom_text(aes(label=paste0(motif_sequences,"/",total_sequences)), angle=90,hjust=-0.5,color="black", size=4)  # Adding text labels

print_plot(STP_subcomplex_bar,"STP_subcomplex_bar.pdf",4,6)

STPxKR_subcomplex_bar<-detected_df2 %>%
    filter(detected==TRUE) %>%
    # create col that says whether seq matches motif or not
    mutate(motif=grepl(".......[ST]P.[KR]....",Sequence)) %>%
    group_by(stage,new_cat) %>%
    summarise(
        total_sequences = n(),
        motif_sequences = sum(motif)
    )  %>%
    mutate(perc_match=motif_sequences/total_sequences*100) %>%
    mutate(stage=factor(stage,levels=c("pro","MI","MII","mitoM"))) %>%
    #select(stage,perc_match) %>% head(10)
    filter(new_cat %in% c("cbf1_3","inner_kt","KMN","dam1")) %>%
    ggplot(aes(x=interaction(stage,new_cat),y=perc_match,fill=new_cat))+
    geom_bar(stat="identity",position="stack")+
    coord_flip()+
    theme_classic()+
    labs(title="[ST]*Px[KR] sites",y="percent_match",x="")+
    scale_y_continuous(expand=expansion(mult=c(0,0.2)))+
    theme(legend.position="none")+
    geom_text(aes(label=paste0(motif_sequences,"/",total_sequences)), hjust=-0.5,color="black", size=4)  # Adding text labels

print_plot(STPxKR_subcomplex_bar,"STPxKR_subcomplex_bar.pdf",6,3)

DENxST_subcomplex_bar<-detected_df2 %>%
    filter(detected==TRUE) %>%
    # create col that says whether seq matches motif or not
    mutate(motif=grepl(".....[DEN].[ST].......",Sequence)) %>%
    group_by(stage,new_cat) %>%
    summarise(
        total_sequences = n(),
        motif_sequences = sum(motif)
    )  %>%
    mutate(perc_match=motif_sequences/total_sequences*100) %>%
    mutate(stage=factor(stage,levels=c("pro","MI","MII","mitoM"))) %>%
    #select(stage,perc_match) %>% head(10)
    filter(new_cat %in% c("cbf1_3","inner_kt","KMN","dam1")) %>%
    # factor levels so that groups go from inner to outer kinetochore
    mutate(new_cat=factor(new_cat,levels= c("cbf1_3","inner_kt","KMN","dam1"))) %>%
    ggplot(aes(x=interaction(stage,new_cat),y=perc_match,fill=new_cat))+
    geom_bar(stat="identity",position="stack")+
    #coord_flip()+
    theme_classic()+
    labs(title="[DEN]x[ST]* sites",y="percent_match",x="")+
    scale_y_continuous(expand=expansion(mult=c(0,0.2)))+
    theme(legend.position="none",axis.text.x = element_text(angle=90,vjust=0.5))+
    geom_text(aes(label=paste0(motif_sequences,"/",total_sequences)), angle=90,hjust=-0.5,color="black", size=4)  # Adding text labels

print_plot(DENxST_subcomplex_bar,"DENxST_subcomplex_bar.pdf",4,6)


DExST_subcomplex_bar<-detected_df2 %>%
    filter(detected==TRUE) %>%
    # create col that says whether seq matches motif or not
    mutate(motif=grepl(".....[DE].[ST].......",Sequence)) %>%
    group_by(stage,new_cat) %>%
    summarise(
        total_sequences = n(),
        motif_sequences = sum(motif)
    )  %>%
    mutate(perc_match=motif_sequences/total_sequences*100) %>%
    mutate(stage=factor(stage,levels=c("pro","MI","MII","mitoM"))) %>%
    #select(stage,perc_match) %>% head(10)
    filter(new_cat %in% c("cbf1_3","inner_kt","KMN","dam1")) %>%
    ggplot(aes(x=interaction(stage,new_cat),y=perc_match,fill=new_cat))+
    geom_bar(stat="identity",position="stack")+
    coord_flip()+
    theme_classic()+
    labs(title="[DE]x[ST]* sites",y="percent_match",x="")+
    scale_y_continuous(expand=expansion(mult=c(0,0.2)))+
    theme(legend.position="none")+
    geom_text(aes(label=paste0(motif_sequences,"/",total_sequences)), hjust=-0.5,color="black", size=4)  # Adding text labels

print_plot(DExST_subcomplex_bar,"DExST_subcomplex_bar.pdf",6,3)

NxST_subcomplex_bar<-detected_df2 %>%
    filter(detected==TRUE) %>%
    # create col that says whether seq matches motif or not
    mutate(motif=grepl(".....[N].[ST].......",Sequence)) %>%
    group_by(stage,new_cat) %>%
    summarise(
        total_sequences = n(),
        motif_sequences = sum(motif)
    )  %>%
    mutate(perc_match=motif_sequences/total_sequences*100) %>%
    mutate(stage=factor(stage,levels=c("pro","MI","MII","mitoM"))) %>%
    #select(stage,perc_match) %>% head(10)
    filter(new_cat %in% c("cbf1_3","inner_kt","KMN","dam1")) %>%
    ggplot(aes(x=interaction(stage,new_cat),y=perc_match,fill=new_cat))+
    geom_bar(stat="identity",position="stack")+
    coord_flip()+
    theme_classic()+
    labs(title="Nx[ST]* sites",y="percent_match",x="")+
    scale_y_continuous(expand=expansion(mult=c(0,0.2)))+
    theme(legend.position="none")+
    geom_text(aes(label=paste0(motif_sequences,"/",total_sequences)), hjust=-0.5,color="black", size=4)  # Adding text labels

print_plot(NxST_subcomplex_bar,"NxST_subcomplex_bar.pdf",6,3)


DENxSTF_subcomplex_bar<-detected_df2 %>%
    filter(detected==TRUE) %>%
    # create col that says whether seq matches motif or not
    mutate(motif=grepl(".....[DEN].[ST]F......",Sequence)) %>%
    group_by(stage,new_cat) %>%
    summarise(
        total_sequences = n(),
        motif_sequences = sum(motif)
    )  %>%
    mutate(perc_match=motif_sequences/total_sequences*100) %>%
    mutate(stage=factor(stage,levels=c("pro","MI","MII","mitoM"))) %>%
    #select(stage,perc_match) %>% head(10)
    filter(new_cat %in% c("cbf1_3","inner_kt","KMN","dam1")) %>%
    ggplot(aes(x=interaction(stage,new_cat),y=perc_match,fill=new_cat))+
    geom_bar(stat="identity",position="stack")+
    coord_flip()+
    theme_classic()+
    labs(title="[DEN]x[ST]*F sites",y="percent_match",x="")+
    scale_y_continuous(expand=expansion(mult=c(0,0.2)))+
    theme(legend.position="none")+
    geom_text(aes(label=paste0(motif_sequences,"/",total_sequences)), hjust=-0.5,color="black", size=4)  # Adding text labels

print_plot(DENxSTF_subcomplex_bar,"DENxSTF_subcomplex_bar.pdf",6,3)

RKxST_subcomplex_bar<-detected_df2 %>%
    filter(detected==TRUE) %>%
    # create col that says whether seq matches motif or not
    mutate(motif=grepl(".....[RK].[ST].......",Sequence)) %>%
    group_by(stage,new_cat) %>%
    summarise(
        total_sequences = n(),
        motif_sequences = sum(motif)
    )  %>%
    mutate(perc_match=motif_sequences/total_sequences*100) %>%
    mutate(stage=factor(stage,levels=c("pro","MI","MII","mitoM"))) %>%
    #select(stage,perc_match) %>% head(10)
    filter(new_cat %in% c("cbf1_3","inner_kt","KMN","dam1")) %>%
    ggplot(aes(x=interaction(stage,new_cat),y=perc_match,fill=new_cat))+
    geom_bar(stat="identity",position="stack")+
    coord_flip()+
    theme_classic()+
    labs(title="[RK]x[ST]* sites",y="percent_match",x="")+
    scale_y_continuous(expand=expansion(mult=c(0,0.2)))+
    theme(legend.position="none")+
    geom_text(aes(label=paste0(motif_sequences,"/",total_sequences)), hjust=-0.5,color="black", size=4)  # Adding text labels

print_plot(RKxST_subcomplex_bar,"RKxST_subcomplex_bar.pdf",6,3)

RKRKxST_subcomplex_bar<-detected_df2 %>%
    filter(detected==TRUE) %>%
    # create col that says whether seq matches motif or not
    mutate(motif=grepl("....[RK][RK].[ST].......",Sequence)) %>%
    group_by(stage,new_cat) %>%
    summarise(
        total_sequences = n(),
        motif_sequences = sum(motif)
    )  %>%
    mutate(perc_match=motif_sequences/total_sequences*100) %>%
    mutate(stage=factor(stage,levels=c("pro","MI","MII","mitoM"))) %>%
    #select(stage,perc_match) %>% head(10)
    filter(new_cat %in% c("cbf1_3","inner_kt","KMN","dam1")) %>%
    ggplot(aes(x=interaction(stage,new_cat),y=perc_match,fill=new_cat))+
    geom_bar(stat="identity",position="stack")+
    coord_flip()+
    theme_classic()+
    labs(title="[RK][RK]x[ST]* sites",y="percent_match",x="")+
    scale_y_continuous(expand=expansion(mult=c(0,0.2)))+
    theme(legend.position="none")+
    geom_text(aes(label=paste0(motif_sequences,"/",total_sequences)), hjust=-0.5,color="black", size=4)  # Adding text labels

print_plot(RKRKxST_subcomplex_bar,"RKRKxST_subcomplex_bar.pdf",6,3)

# PE volcanos ####

# now create category column in the MS df

#monopolin <- c("CSM1","MAM1","HRR25","LRS4")

dep_pe_pg_99_filt_df <-dep_pe_pg_99_filt_df %>% 
    mutate(cat=ifelse(Gene.Names %in% monopolin, "monopolin",
                      ifelse(Gene.Names %in% master_kt_df$Gene,"kinetochore",
                             ifelse(Gene.Names %in% spb$Gene,"spindle pole body",
                                    "other"))))


# factor the levels so they always appear in this order
dep_pe_pg_99_filt_df <-dep_pe_pg_99_filt_df %>% 
    mutate(cat=fct_relevel(cat,c("monopolin","kinetochore","spindle pole body","other")))

# for "dubious" protein names, use yeast systematic names 
# Christos says the Dubious names Wera got from Uniprot
# and I don't find them on SGD so they are not official for yeast

dep_pe_pg_99_filt_df<-dep_pe_pg_99_filt_df %>%
    mutate(name=ifelse(grepl("dubious",name),paste0(Protein,"_",Site),name)) 

# create volcanoes
# only want tag vs tag conditions

# diff stages vs each other
get_volcano_labeled3(dep_pe_pg_99_filt_df,"WT_pro_vs_WT_MI","pe_pro_v_MI_volcano.pdf","pe_pro_v_MI_volcano.html")
get_volcano_labeled3(dep_pe_pg_99_filt_df,"WT_pro_vs_WT_MII","pe_pro_v_MII_volcano.pdf","pe_pro_v_MII_volcano.html")
get_volcano_labeled3(dep_pe_pg_99_filt_df,"WT_pro_vs_WT_mitoM","pe_pro_v_mitoM_volcano.pdf","pe_pro_v_mitoM_volcano.html")
get_volcano_labeled3(dep_pe_pg_99_filt_df,"WT_MI_vs_WT_MII","pe_MI_v_MII_volcano.pdf","pe_MI_v_MII_volcano.html")
get_volcano_labeled3(dep_pe_pg_99_filt_df,"WT_MI_vs_WT_mitoM","pe_MI_v_mitoM_volcano.pdf","pe_MI_v_mitoM_volcano.html")
get_volcano_labeled3(dep_pe_pg_99_filt_df,"WT_MII_vs_WT_mitoM","pe_MII_v_mitoM_volcano.pdf","pe_MII_v_mitoM_volcano.html")

# rank top phospho-sites for each comparison 
# (similar to what I did for protein)

# RANK PHOSPHO FOR go ####

# PHOS new GO section ####

# each stage vs the other 3 stages
# of sigdiff proteins (called by DEP)
# what is median fc for that stage vs other 3 (make sure directions)

pe_diff_cols<-dep_pe_pg_99_filt_df  %>% 
    select(grep("_diff",colnames(.),value=T)) %>% colnames()


# start with pro
grep("pro",pe_diff_cols,value=T)
# one needs to be flipped in direction
# WT_pro_vs_WT_MI_diff
# WT_pro_vs_WT_MII_diff
# WT_pro_vs_WT_mitoM_diff

# of the proteins most diff 
# and which are not KT proteins
# what are these proteins
# create combined list for the 3 comparisons

# always make df with _diffs arranged in same stage order 
# for consistency:
# pro -> MI -> MII -> mitoM
# if you start in the middle of the seq go back to the beginning?

pe_pro_enriched_genes<-bind_rows(dep_pe_pg_99_filt_df  %>%
                                  select(Gene.Names,name,WT_pro_vs_WT_MI_diff) %>%
                                  # the positive values here are proteins higher in pro
                                  arrange(desc(WT_pro_vs_WT_MI_diff)) %>%
                                  #head() # spc42-326,ssz1-477 are most diff higher in prophase vs MI
                                  # filter out kinetochore proteins
                                  #nrow()
                                  filter(!Gene.Names %in% master_kt_df$Gene) %>% #nrow()
                                  # take the top 50 proteins
                                  head(50) %>% select(Gene.Names),
                              dep_pe_pg_99_filt_df  %>%
                                  select(Gene.Names, name,WT_pro_vs_WT_MII_diff) %>%
                                  # the positive values here are proteins higher in pro
                                  arrange(desc(WT_pro_vs_WT_MII_diff)) %>%
                                  #head() # Ndc80-54,Mps1-22 are most diff higher in prophase vs MII
                                  # filter out kinetochore proteins
                                  #nrow()
                                  filter(!Gene.Names %in% master_kt_df$Gene) %>% #nrow()
                                  # take the top 50 proteins
                                  head(50) %>% select(Gene.Names),
                              dep_pe_pg_99_filt_df  %>%
                                  select(Gene.Names,name,WT_pro_vs_WT_mitoM_diff) %>%
                                  # the POSITIVE values here are proteins higher in pro
                                  arrange(desc(WT_pro_vs_WT_mitoM_diff)) %>%
                                  #head() # Mrh1-295,Rsc2-682 are most diff higher in prophase vs mitoM
                                  # filter out kinetochore proteins
                                  #nrow()
                                  filter(!Gene.Names %in% master_kt_df$Gene) %>% #nrow()
                                  # take the top 50 proteins
                                  head(50) %>% select(Gene.Names)) %>%
    distinct() %>%
    unlist() %>% unname()

pe_pro_enriched_result<-gost(pe_pro_enriched_genes,organism="scerevisiae",significant = T,
                          evcodes=T)$result 

pe_pro_enriched_result %>%
    select(term_name,term_id,p_value) %>%
    head(20)
# GO:0006325 chromatin organization
# GO:0006302 double-strand break repair
# GO:0006366 transcription by RNA polymerase II

# copy 3 SITE lists as df to excel
dep_pe_pg_99_filt_df  %>%
    select(Gene.Names,name,WT_pro_vs_WT_MI_diff) %>%
    # the positive values here are proteins higher in pro
    arrange(desc(WT_pro_vs_WT_MI_diff)) %>%
    # filter out kinetochore proteins
    #nrow()
    filter(!Gene.Names %in% master_kt_df$Gene) %>% #nrow()
    # take the top 20 SITES
    head(20) %>% select(name) %>% write_clip()

dep_pe_pg_99_filt_df  %>%
    select(Gene.Names,name,WT_pro_vs_WT_MII_diff) %>%
    # the positive values here are proteins higher in pro
    arrange(desc(WT_pro_vs_WT_MII_diff)) %>%
    # filter out kinetochore proteins
    #nrow()
    filter(!Gene.Names %in% master_kt_df$Gene) %>% #nrow()
    # take the top 20 proteins
    head(20) %>% select(name) %>% write_clip()

dep_pe_pg_99_filt_df %>%
    select(Gene.Names,name,WT_pro_vs_WT_mitoM_diff) %>%
    # the POSITIVE values here are proteins higher in pro
    arrange(desc(WT_pro_vs_WT_mitoM_diff)) %>%
    # filter out kinetochore proteins
    #nrow()
    filter(!Gene.Names %in% master_kt_df$Gene) %>% #nrow()
    # take the top 20 proteins
    head(20) %>% select(name) %>% write_clip()

# MI genes

grep("_MI_",pe_diff_cols,value=T)

pe_MI_enriched_genes<-bind_rows(dep_pe_pg_99_filt_df %>%
                                 select(Gene.Names,name,WT_MI_vs_WT_MII_diff) %>%
                                 # the positive values here are proteins higher in MI vs MII
                                 arrange(desc(WT_MI_vs_WT_MII_diff)) %>%
                                 #head() # SLK19-23, Pes4-58 are most diff higher in MI vs MII
                                 # filter out kinetochore proteins
                                 #nrow()
                                 filter(!Gene.Names %in% master_kt_df$Gene) %>% #nrow()
                                 # take the top 50 proteins
                                 head(50) %>% select(Gene.Names),
                             dep_pe_pg_99_filt_df %>%
                                 select(Gene.Names,name,WT_MI_vs_WT_mitoM_diff) %>%
                                 # the positive values here are proteins higher in MI
                                 arrange(desc(WT_MI_vs_WT_mitoM_diff)) %>%
                                 #head() # MRH1-295,289 are most diff higher in MI vs mitoM
                                 # filter out kinetochore proteins
                                 #nrow()
                                 filter(!Gene.Names %in% master_kt_df$Gene) %>% #nrow()
                                 # take the top 50 proteins
                                 head(50) %>% select(Gene.Names),
                             dep_pe_pg_99_filt_df %>%
                                 select(Gene.Names,name,WT_pro_vs_WT_MI_diff) %>%
                                 # the negative values here are proteins higher in MI
                                 arrange((WT_pro_vs_WT_MI_diff)) %>%
                                 #head() # Isw1-1061,Dsn1-547 are most diff higher in MI vs pro
                                 # filter out kinetochore proteins
                                 #nrow()
                                 filter(!Gene.Names %in% master_kt_df$Gene) %>% #nrow()
                                 # take the top 50 proteins
                                 head(50) %>% select(Gene.Names)) %>%
    distinct() %>%
    unlist() %>% unname()


pe_MI_enriched_result<-gost(pe_MI_enriched_genes,organism="scerevisiae",significant = T,
                         evcodes=T)$result 

pe_MI_enriched_result %>%
    select(term_name,term_id,p_value) %>%
    head(20)
#GO:0022402 cell cycle process
#GO:0006338 chromatin remodeling
#GO:0031023 microtubule organizing center organization
#GO:0010468 regulation of gene expression 
#GO:0051276 chromosome organization

# copy 3 gene lists to excel
dep_pe_pg_99_filt_df%>%
    select(Gene.Names,name,WT_MI_vs_WT_MII_diff) %>%
    # the positive values here are proteins higher in MI vs MII
    arrange(desc(WT_MI_vs_WT_MII_diff)) %>%
    # filter out kinetochore proteins
    #nrow()
    filter(!Gene.Names %in% master_kt_df$Gene) %>% #nrow()
    # take the top 50 proteins
    head(20) %>% select(name) %>% write_clip()

dep_pe_pg_99_filt_df%>%
    select(Gene.Names,name,WT_MI_vs_WT_mitoM_diff) %>%
    # the positive values here are proteins higher in MI
    arrange(desc(WT_MI_vs_WT_mitoM_diff)) %>%
    # filter out kinetochore proteins
    #nrow()
    filter(!Gene.Names %in% master_kt_df$Gene) %>% #nrow()
    # take the top 20 proteins
    head(20) %>% select(name) %>% write_clip()

dep_pe_pg_99_filt_df %>%
    select(Gene.Names,name,WT_pro_vs_WT_MI_diff) %>%
    # the negative values here are proteins higher in MI
    arrange((WT_pro_vs_WT_MI_diff)) %>%
    # filter out kinetochore proteins
    #nrow()
    filter(!Gene.Names %in% master_kt_df$Gene) %>% #nrow()
    # take the top 20 proteins
    head(20) %>% select(name) %>% write_clip()

# MII genes

grep("_MII_",pe_diff_cols,value=T)

pe_MII_enriched_genes<-bind_rows(dep_pe_pg_99_filt_df %>%
                                     select(Gene.Names,name,WT_MI_vs_WT_MII_diff) %>%
                                     # the negative values here are proteins higher in MII
                                     arrange((WT_MI_vs_WT_MII_diff)) %>%
                                     #head() # Spc42-326,Nud1-458 are most diff higher in MII vs MII
                                     # filter out kinetochore proteins
                                     #nrow()
                                     filter(!Gene.Names %in% master_kt_df$Gene) %>% #nrow()
                                     # take the top 50 proteins
                                     head(50) %>% select(Gene.Names),
                                 dep_pe_pg_99_filt_df %>%
                                     select(Gene.Names,name,WT_MII_vs_WT_mitoM_diff) %>%
                                     # the positive values here are sites higher in MII
                                     arrange(desc(WT_MII_vs_WT_mitoM_diff)) %>%
                                     # filter out kinetochore proteins
                                     #nrow()
                                     filter(!Gene.Names %in% master_kt_df$Gene) %>% #nrow()
                                     # take the top 50 proteins
                                     head(50) %>% select(Gene.Names),
                                 dep_pe_pg_99_filt_df %>%
                                     select(Gene.Names,name,WT_pro_vs_WT_MII_diff) %>%
                                     # the negative values here are sites higher in MII
                                     arrange((WT_pro_vs_WT_MII_diff)) %>%
                                     #head() 
                                     # filter out kinetochore proteins
                                     #nrow()
                                     filter(!Gene.Names %in% master_kt_df$Gene) %>% #nrow()
                                     # take the top 50 proteins
                                     head(50) %>% select(Gene.Names)) %>%
    distinct() %>%
    unlist() %>% unname()

pe_MII_enriched_result<-gost(pe_MII_enriched_genes,organism="scerevisiae",significant = T,
                             evcodes=T)$result 

pe_MII_enriched_result %>%
    select(term_name,term_id,p_value) %>%
    head(50)
#GO:0022402 cell cycle process
#GO:0031023 microtubule organizing center organization
#GO:0051300 spindle pole body organization (this was used in the protein plot)
#GO:0000226 microtubule cytoskeleton organization
#sexual reproduction GO:0019953

# copy 3 gene lists to excel
dep_pe_pg_99_filt_df%>%
    select(Gene.Names,name,WT_MI_vs_WT_MII_diff) %>%
    # the negative values here are proteins higher in MII
    arrange((WT_MI_vs_WT_MII_diff)) %>%
    # filter out kinetochore proteins
    #nrow()
    filter(!Gene.Names %in% master_kt_df$Gene) %>% #nrow()
    # take the top 50 proteins
    head(20) %>% select(name) %>% write_clip()

dep_pe_pg_99_filt_df%>%
    select(Gene.Names,name,WT_MII_vs_WT_mitoM_diff) %>%
    # the positive values here are proteins higher in MII
    arrange(desc(WT_MII_vs_WT_mitoM_diff)) %>%
    # filter out kinetochore proteins
    #nrow()
    filter(!Gene.Names %in% master_kt_df$Gene) %>% #nrow()
    # take the top 20 proteins
    head(20) %>% select(name) %>% write_clip()

dep_pe_pg_99_filt_df %>%
    select(Gene.Names,name,WT_pro_vs_WT_MII_diff) %>%
    # the negative values here are proteins higher in MII
    arrange((WT_pro_vs_WT_MII_diff)) %>%
    # filter out kinetochore proteins
    #nrow()
    filter(!Gene.Names %in% master_kt_df$Gene) %>% #nrow()
    # take the top 20 proteins
    head(20) %>% select(name) %>% write_clip()

# mitoM genes

grep("_mitoM_",pe_diff_cols,value=T)

pe_mitoM_enriched_genes<-bind_rows(dep_pe_pg_99_filt_df %>%
                                       select(Gene.Names,name,WT_MI_vs_WT_mitoM_diff) %>%
                                       # the negative values here are sites higher mitoM
                                       arrange((WT_MI_vs_WT_mitoM_diff)) %>%
                                       #head()
                                       # filter out kinetochore proteins
                                       #nrow()
                                       filter(!Gene.Names %in% master_kt_df$Gene) %>% #nrow()
                                       # take the top 50 proteins
                                       head(50) %>% select(Gene.Names),
                                   dep_pe_pg_99_filt_df %>%
                                       select(Gene.Names,name,WT_MII_vs_WT_mitoM_diff) %>%
                                       # the negative values here are sites higher in mitoM
                                       arrange((WT_MII_vs_WT_mitoM_diff)) %>%
                                       # filter out kinetochore proteins
                                       #nrow()
                                       filter(!Gene.Names %in% master_kt_df$Gene) %>% #nrow()
                                       # take the top 50 proteins
                                       head(50) %>% select(Gene.Names),
                                   dep_pe_pg_99_filt_df %>%
                                       select(Gene.Names,name,WT_pro_vs_WT_mitoM_diff) %>%
                                       # the negative values here are sites higher in mitoM
                                       arrange((WT_pro_vs_WT_mitoM_diff)) %>%
                                       # filter out kinetochore proteins
                                       #nrow()
                                       filter(!Gene.Names %in% master_kt_df$Gene) %>% #nrow()
                                       # take the top 50 proteins
                                       head(50) %>% select(Gene.Names)) %>%
    distinct() %>%
    unlist() %>% unname()

pe_mitoM_enriched_result<-gost(pe_mitoM_enriched_genes,organism="scerevisiae",significant = T,
                               evcodes=T)$result 

pe_mitoM_enriched_result %>%
    select(term_name,term_id,p_value) %>%
    head(50)

#GO:0006997 nucleus organization
#GO:0060260 regulation of transcription initiation by RNA polymerase II
#GO:0022402 cell cycle process
#GO:0031023 microtubule organizing center organization
#GO:0051300 spindle pole body organization (this was used in the protein plot)
#GO:0000226 microtubule cytoskeleton organization
#

# copy 3 gene lists to excel
dep_pe_pg_99_filt_df %>%
    select(Gene.Names,name,WT_MI_vs_WT_mitoM_diff) %>%
    # the negative values here are sites higher mitoM
    arrange((WT_MI_vs_WT_mitoM_diff)) %>%
    # filter out kinetochore proteins
    #nrow()
    filter(!Gene.Names %in% master_kt_df$Gene) %>% #nrow()
    # take the top 50 proteins
    head(20) %>% select(name) %>% write_clip()

dep_pe_pg_99_filt_df %>%
    select(Gene.Names,name,WT_MII_vs_WT_mitoM_diff) %>%
    # the negative values here are sites higher in MII
    arrange((WT_MII_vs_WT_mitoM_diff)) %>%
    # filter out kinetochore proteins
    #nrow()
    filter(!Gene.Names %in% master_kt_df$Gene) %>% #nrow()
    # take the top 20 proteins
    head(20) %>% select(name) %>% write_clip()

dep_pe_pg_99_filt_df %>%
    #filter(Gene.Names=="dubious205")
    select(Gene.Names,name,WT_pro_vs_WT_mitoM_diff) %>%
    # the negative values here are proteins higher in mitoM
    arrange((WT_pro_vs_WT_mitoM_diff)) %>%
    # filter out kinetochore proteins
    #nrow()
    filter(!Gene.Names %in% master_kt_df$Gene) %>% #nrow()
    # take the top 20 proteins
    head(20) %>% select(name) %>% write_clip()

## phospho combi GO bar ####

# create combined p-value plot of 12 enriched terms.
# bar plot
# y axis is the terms
# x axis is the p-value (-log)
# can draw a line for 0.05

#pro terms
## GO:0006325 chromatin organization
# GO:0006302 double-strand break repair
# GO:0006366 transcription by RNA polymerase II
# MI terms
# #GO:0022402 cell cycle process
#GO:0006338 chromatin remodeling
#GO:0031023 microtubule organizing center organization
#GO:0010468 regulation of gene expression 
#GO:0051276 chromosome organization
# MII terms
# #GO:0022402 cell cycle process
#GO:0031023 microtubule organizing center organization
#GO:0051300 spindle pole body organization (this was used in the protein plot)
#GO:0000226 microtubule cytoskeleton organization
#sexual reproduction GO:0019953
# mitoM terms
#GO:0006997 nucleus organization
#GO:0060260 regulation of transcription initiation by RNA polymerase II
#GO:0022402 cell cycle process
#GO:0031023 microtubule organizing center organization
#GO:0051300 spindle pole body organization (this was used in the protein plot)
#GO:0000226 microtubule cytoskeleton organization
#

pe_enriched_df<-rbind(
    pe_pro_enriched_result %>%
        select(term_name,term_id,p_value) %>% 
        filter(term_id %in% c("GO:0006325","GO:0006302","GO:0006366")) %>%
        cbind(stage="pro"),
    
    pe_MI_enriched_result %>%
        select(term_name,term_id,p_value) %>% 
        filter(term_id %in% c("GO:0006338","GO:0051276","GO:0010468")) %>%
        cbind(stage="MI"),
    
    pe_MII_enriched_result %>%
        select(term_name,term_id,p_value) %>% 
        filter(term_id %in% c("GO:0022402","GO:0051300","GO:0019953","GO:0043934")) %>%
        cbind(stage="MII"),
    
    pe_mitoM_enriched_result %>%
        select(term_name,term_id,p_value) %>% 
        filter(term_id %in% c("GO:0006997","GO:0000226","GO:0060260")) %>%
        cbind(stage="mitoM")) 

pe_combi_GO_bar<-pe_enriched_df %>%
    mutate(term=paste0(term_name," ",term_id)) %>%
    mutate(stage=factor(stage,levels=c("pro","MI","MII","mitoM"))) %>%
    mutate(log_pval = -log10(p_value)) %>%
    group_by(stage) %>%
    # Re-factor term_name within each stage so only 3 terms show up per facet
    mutate(term = factor(term, levels = term[order(log_pval)])) %>%
    ggplot(aes(x = log_pval, y = term, fill = stage)) +
    geom_bar(stat = "identity") +
    facet_wrap(~stage, scales = "free_y",ncol=1) +
    labs(
        x = expression(-log[10](p~value)),
        y = "",
        title = ""
    ) +
    theme_minimal() +
    theme(strip.text = element_text(face = "bold"),
          legend.position="none")+
    geom_vline(xintercept=-log10(0.05),linetype="dashed")

print_plot(pe_combi_GO_bar,"pe_combi_GO_bar.pdf",8,4)

# export dfs ####

saveRDS(dsn1_dep_df,"dsn1_dep_df.RDS")
saveRDS(dep_pg_df,"dep_pg_df.RDS")
saveRDS(dep_pe_pg_99_filt_df,"dep_pe_pg_99_filt_df.RDS")
```
